# Sequential Penta-Omic Extraction Method Using Single Biospecimens of Post-mortem Human Brain

**DOI:** 10.64898/2026.06.26.734872

**Authors:** Scott P. Lyons, Brandie M. Ehrmann, Thomas S. Webb, Cristina Arciniega, Laura E. Herring, Shengru Guo, Stuart Parnham, William K. Scott, Piotr A. Mieczkowski, Jeffrey M. Macdonald

**Author notes:** Co-senior authors contributed equally (Piotr A. Mieczkowski, Jeffrey M. Macdonald).

## Abstract

A multi-omic approach utilizing a single biospecimen is important to avoid intra-sample heterogeneity associated with testing multiple omic single-samples, and for more efficient use of small volumes of precious biopsies (<30 mg). This is especially true for the microanatomy of post-mortem human brain samples. Using post-mortem human brain biospecimens from the NIH NeuroBioBank, a penta-omic sequential extraction method is described, *Si*multaneous *M*etabolomic, *P*roteomic, *L*ipidomic – *D*NA, *R*NA *Ex*traction (SiMPL-DREx). Each sequential omic extract was compared to those obtained by the gold standard single omic method. Preserving RIN is critical for brain and tissue banks, as it is a primary measure of tissue quality. For all five omic extracts, the tissue integrity numbers and omic profiles did not significantly differ from those obtained by the respective omic gold standard method. Unlike past multi-omic studies, this study quantified the relative solvent percentages and upstream losses for both the organic and aqueous phases, confirming an omics loss of under 5%.

## INTRODUCTION

A single-biospecimen multi-omic extraction strategy is essential to minimize intra-sample heterogeneity, to enable efficient use of limited biopsy material, and to preserve highly requested human brain tissues used in neurodegenerative research. Sequential extraction from a single specimen is particularly important for validating emerging micro- and single-cell multi-omic imaging approaches. Existing multi-omic workflows include a quad-omic sequential extraction developed for archaeological tissue¹ and established tri-omic biphasic methods based on Folch (MPLEx)^2,3^ or Matyash^4^ (SiMPLEx)^5–9^ chemistries; however, these approaches exclude RNA from the workflow.

Early attempts to extend MPLEx to penta-omic extraction revealed partial degradation of RNA, as measured by RNA integrity number (RIN), in the post-extraction pellet relative to direct tissue extraction using TRIzol.^10,11^ Chaotropic agents commonly used for RNA isolation, including TRIzol™ and RNeasy^®^ reagents, can hydrolyze lipids and compromise metabolomic analyses by introducing NMR spectral artifacts and affecting LC–MS ionization efficiency, background signal, and equilibration time.^11^ To improve RNA preservation, a more recent penta-omic Matyash workflow incorporated RNAlater^®^—an antichaotropic reagent containing ammonium sulfate, citrate, and EDTA—prior to extraction.^12^ When monophasic and biphasic methods were compared across six human colon biopsies, the SiMPLEx-based workflow best preserved RIN; however, RIN values were slightly reduced compared to the gold standard RNA extraction method.^12^ Although RNAlater improves RNA preservation, it will affect LC-MS ionization and adduct formation and taint NMR metabolomic analyses. Previous SiMPLEx studies did not extract RNA and determine RIN values, so it unknown whether RIN is preserved without RNAlater^®^. Most existing multi-omic workflows estimate extraction efficiency by comparing omic profiles and internal standards to single-omic gold standard methods^1,3,5,7–9,12^, however, upstream losses introduced at each extraction step are typically not quantified or retained for potential recovery^13^. Together, these limitations highlight the need for a gentler, fully integrated penta-omic extraction strategy that preserves the integrity of all molecular classes while explicitly accounting for upstream extraction-associated losses.

Maintaining RIN values in sequential extractions is especially important for human brain samples from repositories such as the NIH NeuroBioBank (NBB), because RIN is a primary tissue quality control (QC) metric and is measured at a specific anatomical site (the occipital and frontal poles). As a result, repeated requests for small, high-demand regions such as hippocampus or nucleus accumbens preferentially deplete samples with RIN >7, leaving behind tissues with lower RIN values—even though RIN may not reflect the integrity of other omic extracts. Before evaluating whether RIN correlates with non-transcriptomic data quality, a single-biospecimen penta-omic extraction method that preserves RNA integrity while defining upstream workflow losses is required.

Here we present SiMPL-DREx (Simultaneous Metabolomic, Proteomic, Lipidomic–DNA and RNA Extraction), a sequential penta-omic extraction method derived from the MTBE-based biphasic SiMPLEx workflow.^1,5–9,12^ SiMPL-DREx preserves RNA integrity by minimizing RNA residence time in the aqueous phase while maintaining low-temperature conditions (∼4 °C) throughout to suppress enzymatic activity. SiMPL-DREx has more rapid cell disruption and lipid extraction steps and a single biphasic extraction step compared to previous SiMPLEx workflows^5–9,12^ to reduce RNA exposure to RNases and accelerate throughput. Using 25 mg of postmortem human brain tissue from five donors, we validate SiMPL-DREx by benchmarking each omic layer against its respective single-omic gold standard extraction, quantifying extraction-associated losses, and demonstrating preservation of RIN alongside high-quality metabolomic, lipidomic, proteomic, and genomic profiles.

## RESULTS

### Determination of Solvent Ratios in Organic and Aqueous Phase of SiMPL-DREx

The solvent ratios of organic and aqueous phases in SiMPLEx have never been reported.^1,5–9,12^ It is important to determine this ratio because RNase and RNA in MeOH:water mixtures dictating the RNA/RNase resident time thereby affecting RIN values. **Table 1** lists the relative ratios of MTBE:MeOH:H_2_O in the organic and aqueous phases at four different MTBE:MeOH:H_2_O initial extraction ratios. At a 1:1:1 MTBE:MeOH:H_2_O initial extraction ratio there is no phase separation visible. Most interesting is that methanol remains about 35% in the aqueous phase regardless of the initial MTBE:MeOH:H_2_O extraction ratios.

**Table 1:** MTBE:MeOH:H2O extraction ratios and the same in the aqueous and organic phases.

| Conditions | MTBE:MeOH:H<sub>2</sub>O Ratios |  |  |  |
| --- | --- | --- | --- | --- |
|  | % Volumes | Vol Ratios | Organic Layer | Aqueous Layer |
| Pure Solvents | 62:21:17 | 900:300:250 | 76±1.0 : 18±0.1 : 6±1.0 | 21±0.5 : 36±0.3 : 44±0.3 |
| Pure Solvents | 60:20:20 | 750:250:250 | 76±0.3 : 15±0.1 : 9±0.3 | 14±0.1 : 34±0.1 : 52±0.2 |
| Pure Solvents | 56:24:20 | 700:300:250 | 67±0.6 : 26±0.2 : 12±0.7 | 25±0.5 : 35±0.4 : 40±0.8 |
| Pure Solvents | 53:27:20 | 667:333:250 | 57 : 35 : 18 | 31 : 35 : 33 |
| Pure Solvents | 33:33:33 | 330:330:330 | No Phase separation | No Phase separation |

### SiMPL-DREx Workflow and Comparison to the Respective Single Omic Extraction Method

The demographics, RIN values, and postmortem interval (PMI) for each of the 5 donor samples are given in the table in **Supplementary Table 1A**. The SiMPL-DREx workflow is illustrated in **Figure 1A**. Briefly, ∼25 mg of tissue is extracted using methyl *tert*-butyl ether (MTBE) and methanol (MeOH), with phase separation induced by the addition of water (H₂O). The upper organic phase contains lipids, while the lower aqueous phase contains metabolites; these were compared to their respective gold-standard extraction methods, Matyash^4^ and 80% MeOH^14^ as performed in previous multi-omic sequential extraction method studies^1,5,7–12^ (**Fig. 1A**). Approximately 5 mg of the resulting pellet is used for RNA and DNA extraction via Zymo- Quick columns, and the remaining ∼13 mg is used for global proteomic analysis. Both were compared to extractions performed on tissue from the same donor using the corresponding single-omic methods (**Fig. 1A**).

**Figure 1:**
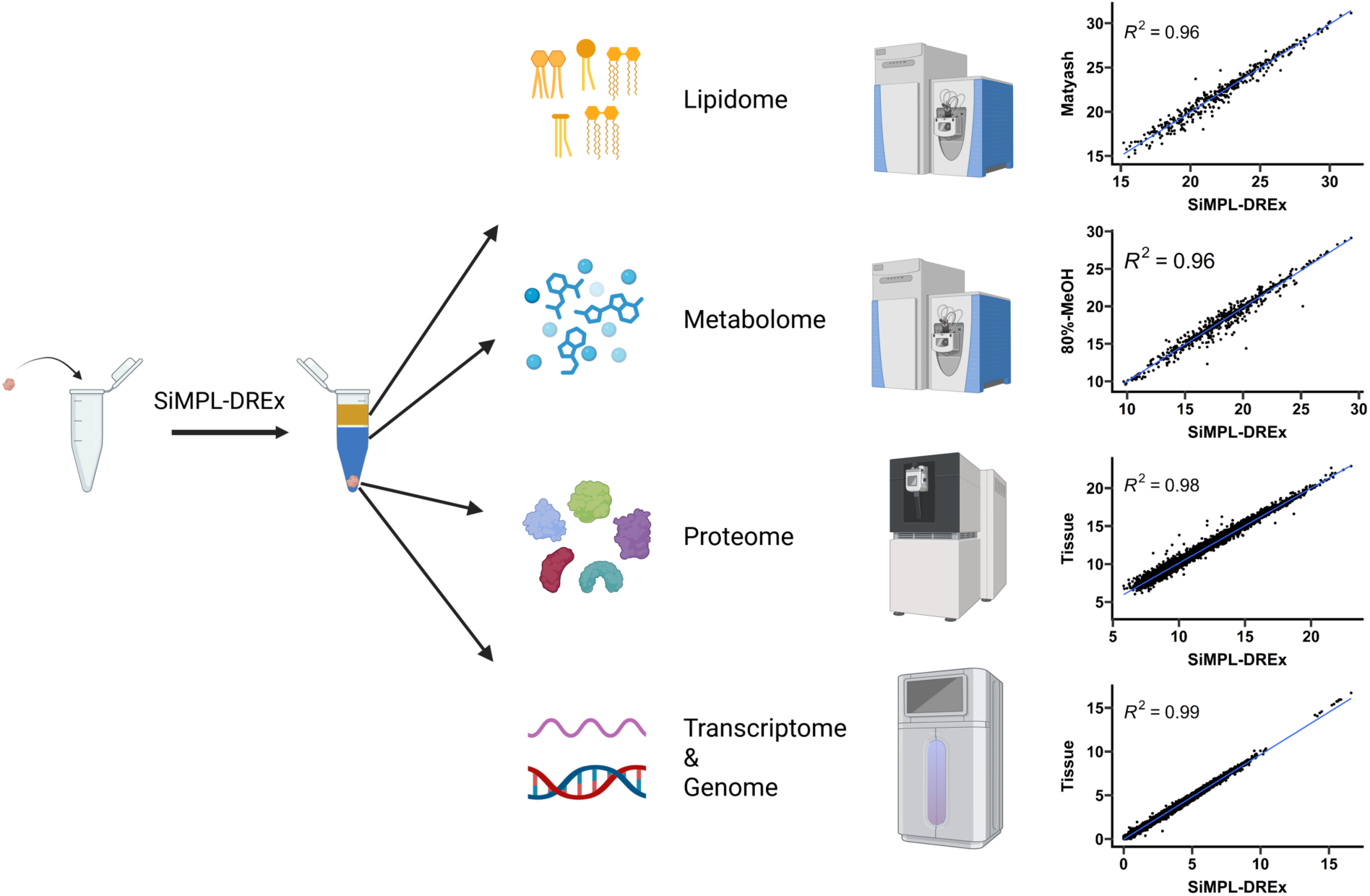
(**A**) **Workflow.** A single brain biopsy specimen is either extracted using the SiMPL-DREx method or the standard method for each extraction method. The panel lists the demographics and key sample information. (**B**) **Correlation plots** on the right represent the correlation coefficients for the top-to-bottom order, namely the lipidome, metabolome, proteome, and transcriptome.

To assess method reproducibility, the intensity of each molecular feature in the lipidomic, metabolomic, proteomic, and total RNA-seq datasets from each of the five donors was plotted against the corresponding feature obtained using the respective gold-standard methods (**Fig. 1B**). All comparisons yielded coefficients of determination (R²) greater than 0.96, demonstrating the robustness of the SiMPL-DREx workflow relative to single-omic extraction approaches. To identify potential biases within each omic dataset, upstream extraction losses were examined to determine whether they correlated with molecular features exhibiting greater than two-fold differences in abundance.

### RNA and DNA Extracts (SiMPL-DREx pellet versus whole tissue)

For the RNA and DNA extracts, the primary omic quality control (QC) factor is the RIN and DIN value, respectively. The individual RIN values of each donor were nearly identical between the two methods using Zymo-Quick solid-phase extraction (**Fig.2A**). To expand the cohort, eight additional donors were selected using demographic and comorbidity criteria matching the five initial samples subjected to multi-omic analysis (**Fig. 1A**; **Supplementary Table 1A**). A paired t-test comparing the extraction methods revealed no significant within-individual differences in RIN values across the 13 donors (*p* = 0.26; **Fig. 2A**, **Supplementary Table 1B**). Notably, even the most degraded sample (RIN = 1.0) maintained a DV200 of 42, safely exceeding the established quality threshold of 30 required for viable short-read RNA-seq performance (**Supplementary Table 1B** and **1C).** In contrast, the DIN was significantly lower for each donor subjected to SiMPL-DREx extraction (*p* = 0.007; **Fig.2A**) possibly due to the increased fragmentation caused by vortexing during the sequential lipidomic and metabolomic extractions. **Figure 2B** is the PCA plot of the two groups showing only minor transcriptional differences within individuals by extraction method, and some separation of individuals regardless of extraction method. Specifically, donors 1 and 4 cluster separately from the remaining samples in PC1, and from each other in PC2, which is also shown in the heatmap (**Fig. 2C**). indicating substantial transcriptomic divergence specific to those two samples. Interestingly, donors 1 and 4 (males) both have relatively higher RIN values compared to donors 2, 3, and 5 (females); therefore, RIN is confounded by sex in those two groups of donors, and it cannot be determined whether the clustering is due to differences in RNA integrity or sex (**Fig. 2A**). Also, considering all 13 donors, no correlation (r=0.004, p=0.99) is observed between PMI and the initial RIN values recorded at the time of sample acquisition (**Fig. 1A**; **Supplementary Table 1A**). Paired statistical analysis of ∼19,000 transcripts revealed no transcript with significant (adjusted p-value < 0.05) 2-fold difference between SiMPL-DREx pellet and whole-tissue extracts (**Fig. 2D**). The correlation of variation was also similar (**Fig. 2E**). The lack of difference in extracted transcripts between methods is reflected by a correlation of determination (R^2^) value of 0.99 (**Fig. 1B**). Also supporting the strong correlation in **Figure 1B** is that no RNA is detected in the upstream lipidome and metabolome extracts (**Supplementary Table 1D**). Similar to RNA, no DNA was detected in the upstream lipidome and metabolome extracts (**Supplementary Table 1D**).

**Figure 2:**
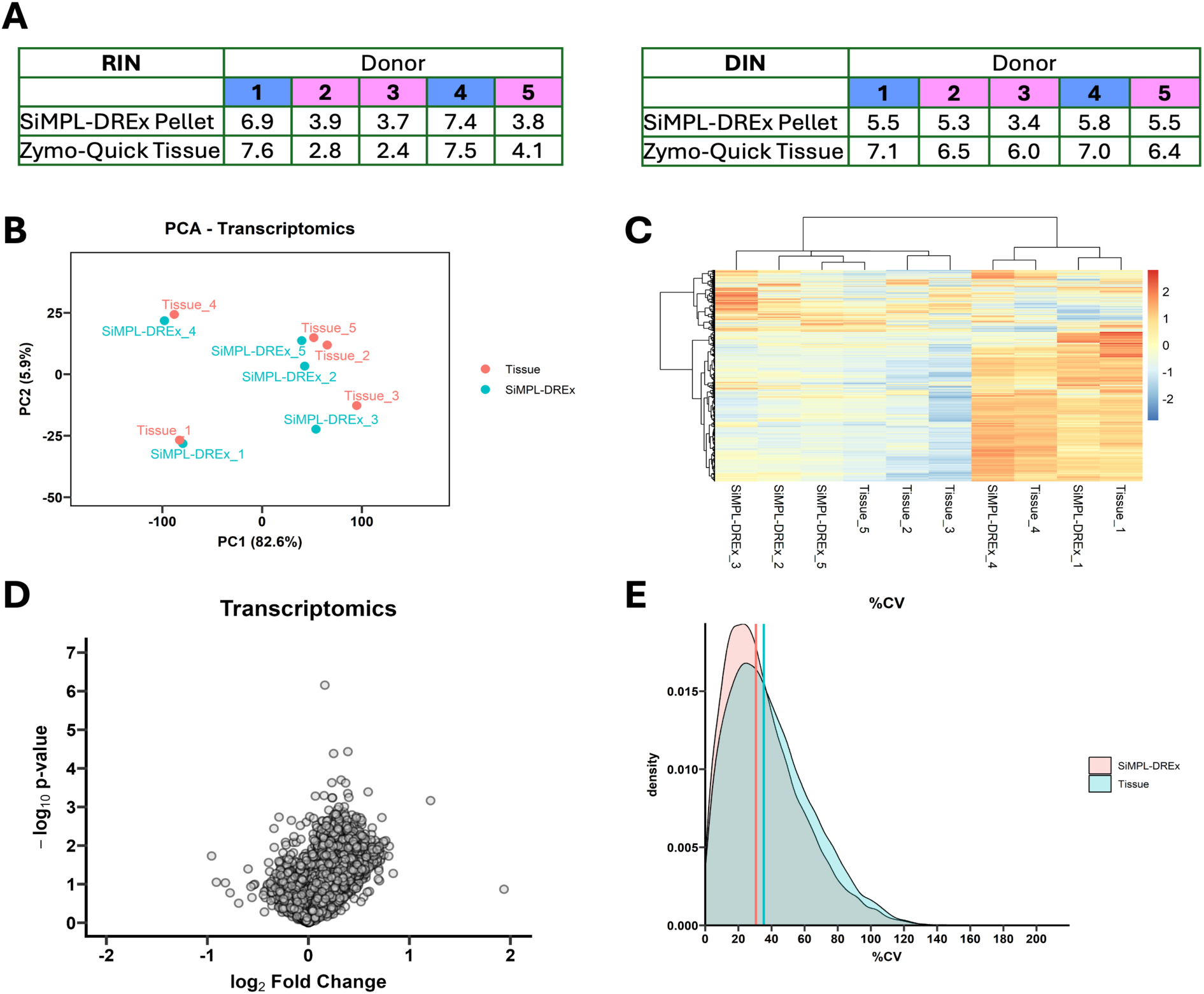
DNA/RNA extracts. **(A)** Comparison of the RNA Integrity Number (RIN) and DNA Integrity Number (DIN) between patients by extraction technique. The patient numbers colored blue represent male samples and pink represent females. **(B)** Principal Component Analysis (PCA) of the transcriptomic data from 5 control and 5 SiMPL-DREx-extracted samples. **(C)** Heatmap of all samples, clustering based on the Ward.D2 method. **(D)** Volcano plot showing the differential expression between SiMPL-DREx and Tissue control methods. Transcripts with an adjusted p-value < 0.05 & log2FC > 0.5 are colored red, those with an adjusted p-value < 0.05 and log2FC < -0.5 are colored blue out of a total of 19,054 transcripts**. (E)** Coefficient of variation of both conditions. The vertical line indicates the median CV.

### Proteomic Extracts (SiMPL-DREx pellet vs whole tissue)

**Figure 3A** shows no significant differences in Protein Integrity Numbers (PIN)^15^ across donors, extraction methods, sex, or RIN. Principal component analysis (**Fig. 3B**) of five donors extracted by both methods demonstrates primary separation by extraction method, with a consistent downward-left shift for SiMPL-DREx relative to whole tissue protein extraction. However, paired statistical analysis of ∼9,000 quantified proteins (**Fig. 3C**) revealed no proteins with significant (adjusted p-value < 0.05) 2-fold difference between SiMPL-DREx pellet and whole-tissue extracts. **Figure 3D** presents protein-level coefficients of variation (%CV; SD/mean) as a measure of reproducibility. In label-free global proteomics, median %CV <20% is generally acceptable. SiMPL-DREx pellet and whole-tissue extracts exhibited median %CVs of 17.19% and 20.46%, respectively (**Fig. 3D**). Because both extraction methods were applied to the same donor samples, biological variation is equivalent, and technical reproducibility is expected to be comparable. Thus, differences in %CV distributions (**Fig. 3D**) could reflect extraction-related bias caused by upstream sample loss.

**Figure 3:**
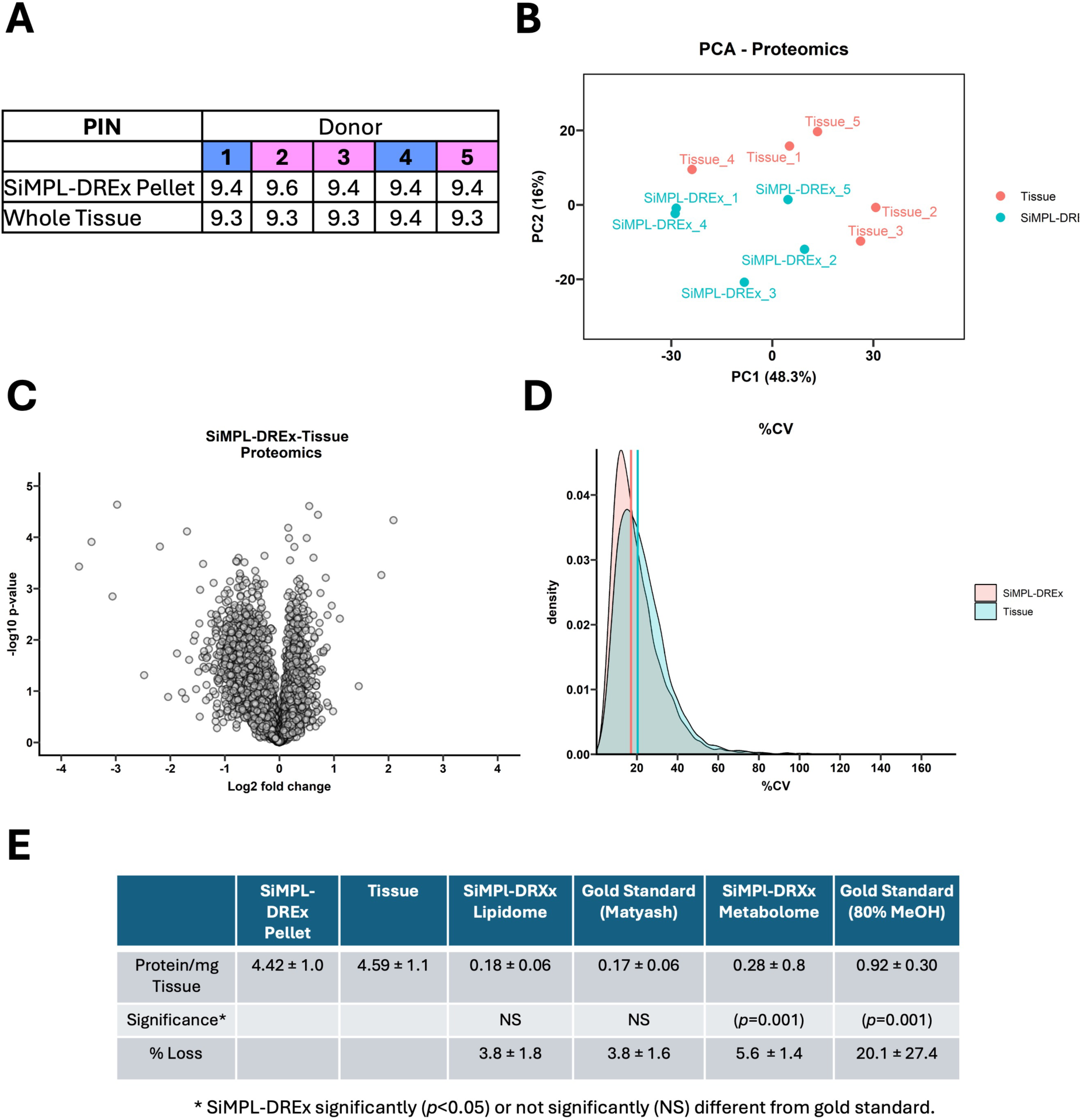
Protein extracts. (**A**) Comparison of the Protein Integrity Number (PIN) between patients and by extraction method. The patient numbers colored blue represent male samples and pink represent females. (**B**) Principal Component Analysis (PCA) of the proteomics data from 5 control and 5 SiMPL-DREx-extracted samples. (**C**) Volcano plot showing the differential expression between SiMPL-DREx and Tissue control methods. Proteins with an adjusted p-value < 0.05 & log2FC > 0.5 are colored red, those with an adjusted p-value < 0.05 & log2FC < -0.5 are colored blue. (**D**) The coefficient of variation (CV) of the SiMPLE-DREx and Tissue samples. (**E**) The upstream protein loss in SiMPL-DREx and gold standards.

To determine if the slight downward shift in %CV distribution of SiMPL-DREx (**Fig. 3D**) could be attributed to upstream losses, protein content in the lipidomic and metabolomic extracts was quantified by BCA assay (Fig. 3E). Total protein per mg tissue did not differ significantly between SiMPL-DREx pellet and whole-tissue extracts. However, small proportions of protein were detected in the SiMPL-DREx lipidome (3.8 ± 1.8%; 0.18 μg/4.59 μg total protein) and metabolome (5.6 ± 1.41%; 0.28 μg/4.59 μg total protein) fractions, suggesting upstream protein loss. This may contribute to the altered %CV distribution (**Fig. 3D**). However, the SiMPL-DREx protein loss in the metabolomic fraction is 5.6 ± 1.4% or about four-fold less than the 80% MeOH metabolomic gold-standard extract which contained 20.8 ± 7.6% (**Fig. 3D**). To determine whether the lost protein represented a subset of tissue proteins or distinct species, global proteomics was performed on the 80% MeOH extract. **Supplementary Figure 1A** is a Venn diagram of the complementary and different proteins extracted between the tissue control and 80% MeOH. There are 9,002 and 7,322 proteins identified in the SiMPL-DREx and tissue versus the 80% MeOH, respectively, with 7,007 proteins in common. Comparative analysis of physicochemical propertie^11^ revealed that proteins uniquely identified following the 80% methanol extraction exhibited a rightward shift in hydrophobicity relative to those uniquely identified using urea-based lysis, indicating enrichment of hydrophobic proteins (**Supplementary Fig. 1A**). In parallel, the isoelectric point (pI) distribution showed a modest shift toward higher pI values in the methanol-extracted proteome (**Supplementary Fig. 1B**). On the other hand, the molecular weight distribution of the proteins showed no distinction between the two conditions (**Supplementary Fig. 1C**). Together, these data suggest that methanol extraction preferentially recovers a physicochemically distinct subset of proteins characterized by increased hydrophobicity and a tendency toward more basic amino acid composition.

### Metabolomic Extracts (SiMPL-DREx vs 80% MeOH)

After transferring the upper lipid fraction, 800 μL of MeOH is added to the lower phase comprised of 35% MeOH (**Table** 1) to achieve ∼80% MeOH.^5^ Then the pellet is resuspended, vortexed for 10s, and centrifuged to further precipitate protein, especially RNase (**Supplementary Fig. 2**). The metabolomic extracts from SiMPL-DREx and 80% MeOH of whole-tissue yielded a coefficient of determination (R²) of 0.96 indicating similar extraction efficiencies between the two methods for all metabolites detected by untargeted LC-MS/MS metabolomics (**Fig. 1B**). The PCA plot shows the distribution of the 5 donors extracted by both methods (**Fig. 4A**). As expected for similar solvents schemes in both methods, donors 1-5 cluster with one another more than they cluster by extraction methods supporting the R² value of 0.96, suggesting little difference in metabolomic profile by extraction method. Interestingly, the clustering and distribution of the respective donors are similar to those from the transcriptomic data (**Fig. 2B**), wherein donors 1 and 4 cluster separately from the remaining samples in PC1 (accounting for 53.1% of variance), and from each other in PC2 (accounting for 15.5% of variance) indicating substantial metabolomic divergence specific to those two samples. (**Fig. 4A**). The clustered heatmap shows the first division of samples grouping male donors 1 and 4 with each other and separated from female donors 2, 3, and 5 (**Fig. 4B**). The volcano plot shows that SiMPL-DREx extracts 13 metabolites at significantly lower levels (adjusted p-value < 0.05 & ≥2-fold decrease) and 12 metabolites at significantly higher levels (adjusted p-value < 0.05 & ≥2-fold increase) out of 653 metabolites (**Fig. 4C**). **Figure 4D** shows the coefficients of variation for both conditions with the vertical line indicating the median CV. The %CV profiles are the similar but shifted slightly higher for the 80% MeOH, suggesting that there is little difference between methods. There is a relatively high median CV of 40 and 44%, which could indicate donor-specific biological and environmental variations between the donors. To evaluate if metabolite differences between the two methods (**Fig. 4C**) could be due to losses in the upstream SiMPL-DREx lipidomic extraction step, **Figure 4E** displays the percentage loss of specific metabolites in the upper lipidome fraction. Metabolites were analyzed in the lipidomic fraction by ^1^H NMR spectroscopy. To determine if the loss would affect the overall - omic profile by preferentially extracting nonpolar amino acids, the percent loss was correlated to the partition coefficients (LogP) of detected amino acids and the hydrophobic parameters (π) of their side chains^16^ in **Figure 4F**. There is a correlation (R^2^=0.82) of LogP (**Fig. 4F**, top graph) and π (**Fig. 4F**, bottom graph) with loss of more nonpolar amino acids in the lipid fraction.

**Figure 4:**
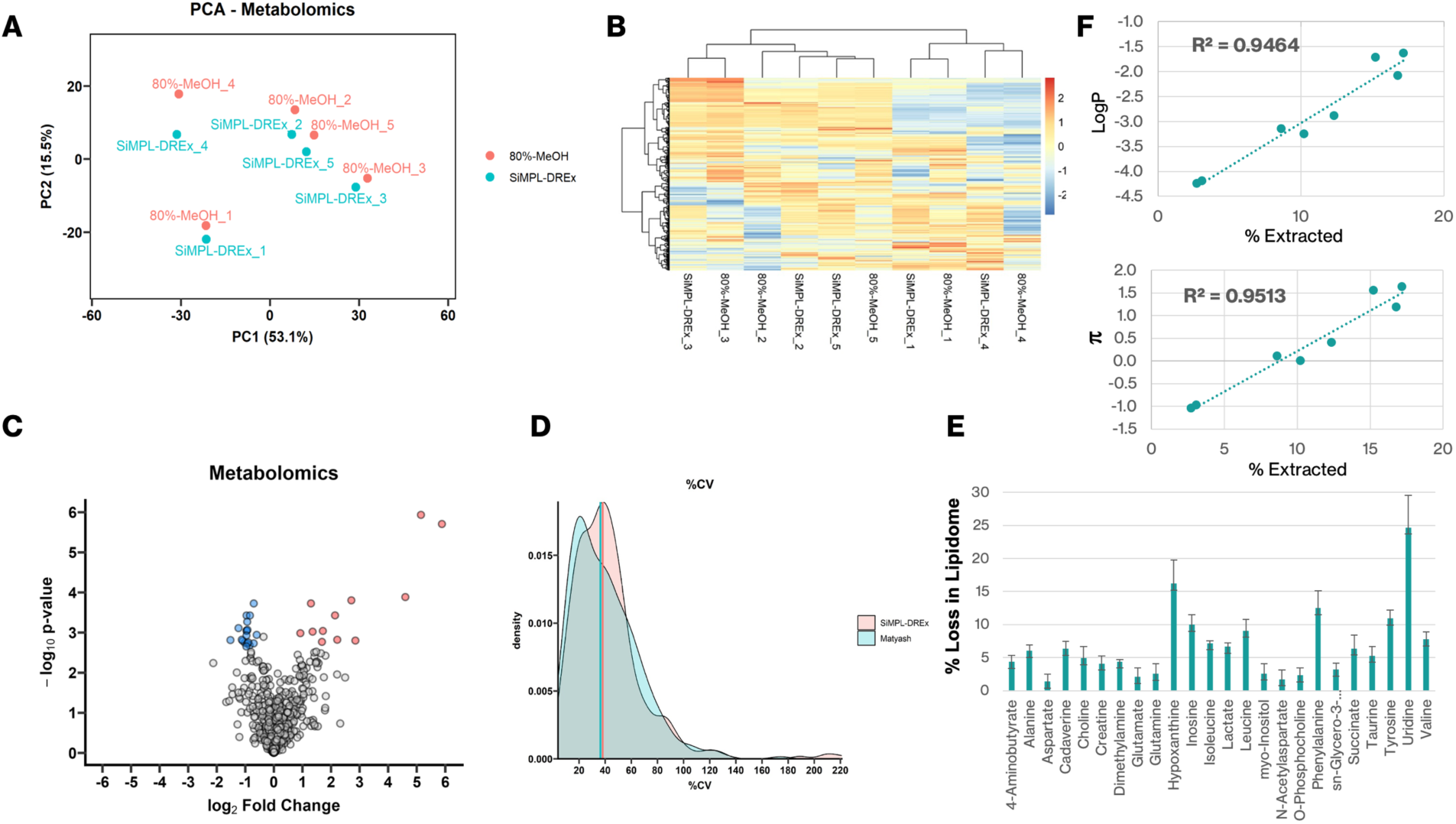
(**A)** Principal Component Analysis (PCA) of the metabolomics data from 5 control and 5 SiMPL-DREx-extracted samples. (**B**) Heatmap of all samples, clustering based on the Ward.D2 method. (**C**) Volcano plot showing the differential metabolite levels between SiMPL-DREx and the 80% MeOH gold standard methods. Metabolites with an adjusted p-value < 0.05 and a log2FC > 0.5 are colored in red, those with a log2FC < -0.5 are colored blue. (**D**) Coefficient of variation of both conditions. The vertical line indicates the median CV. (**E**), upstream metabolomic losses in the SiMPL-DREx lipidomic extract and (**F**) the correlation of percent loss of amino to the hydrophobic parameters, octanol-water partitioning coefficient (LogP) and Hansch pi (π).

### Lipidomic Extracts (SiMPL-DREx vs Matyash)

There were the same 391 lipids identified in both the SiMPL-DREx and Matyash^4^ extraction methods. **Figure 5A** is the PCA plot of SiMPL-DREx versus Matyash showing donors 1-5 for each extraction. The donors 1-5 cluster better with one another than they do with the two methods, and within donors there is no clear separation of donors 1 and 4 from donors 2, 3, and 5. Surprisingly, the clustered heatmap in **Figure 5B** has a large branch distance for the dendrogram connecting donor 3 lipid profiles as well as donor 2. Even though they cluster in the PCA score plot (**Fig. 5A**) the heatmap indicates high dissimilarity. PC1 and PC2 together only account for 39.9% of the variance (**Fig. 5A**), while PC3 and PC4 account for 15.2% and 12.5% or together 27.7% of the variance (**Supplementary Fig. 3A**). In addition, PC2 versus PC3 and PC3 versus PC4 plots both separate by method but PC3 and PC4 have the largest separation between donor 3 and 2, respectively, (**Supplementary Fig. 3A**) supporting the divergence of these donors in the clustered heatmap (**Fig. 5B**). The volcano plot of SiMPL-DREx versus Matyash revealed no lipids with significant (adjusted p-value < 0.05) 2-fold difference between SiMPL-DREx pellet and whole-tissue extracts (**Fig. 5C**). **Figure 5D** represents lipid-level coefficients of variation (%CV; SD/mean) as a measure of reproducibility. The median CVs are nearly identical for tissue and SiMPL-DREx extracts at 38% and 39%, respectively. In short, SiMPL-DREx shows very little difference in extracted lipids, likely because they are essentially the same procedure of the same tissues (compare workflows in **Supplementary Figs. 2** and **4**).

**Figure 5:**
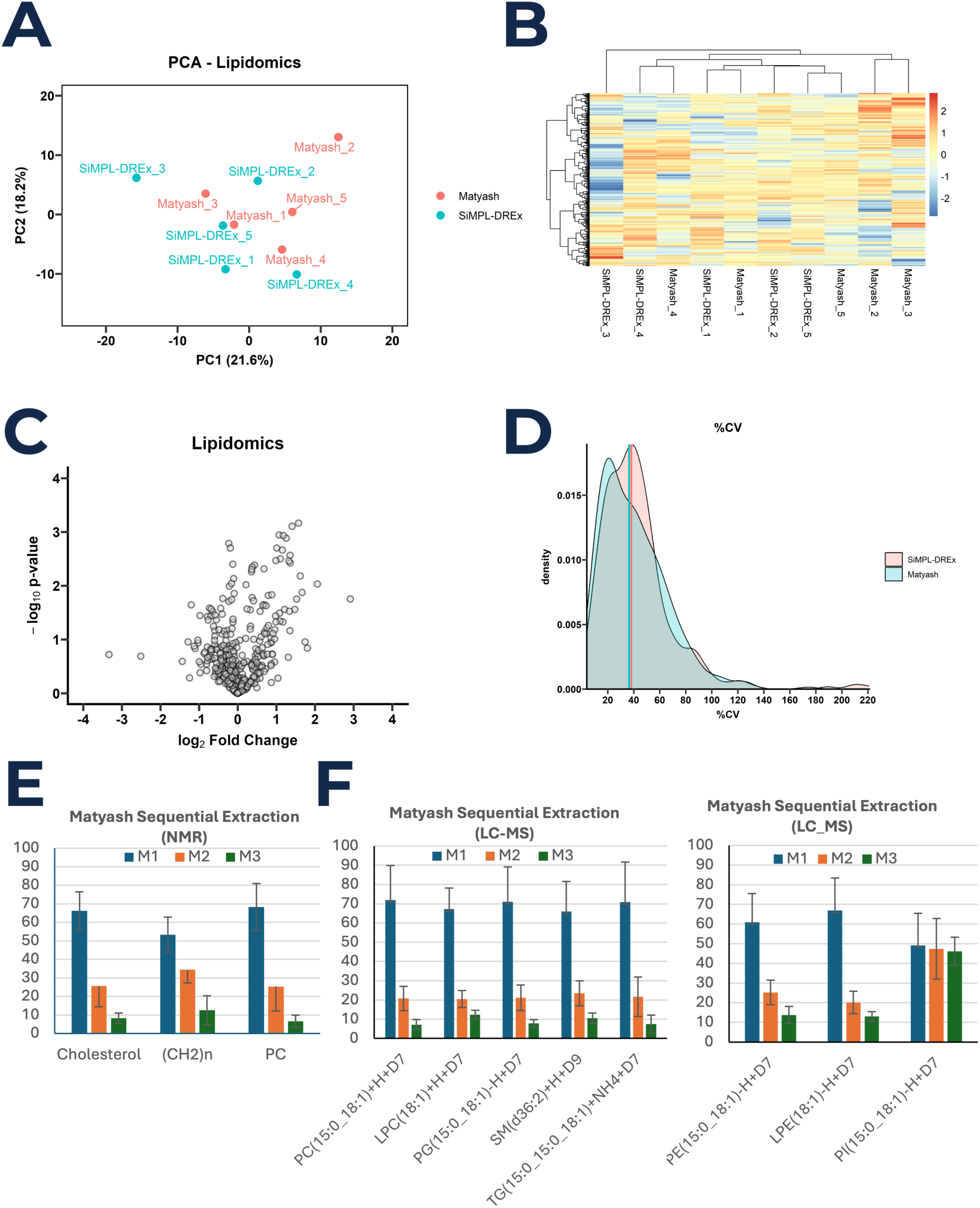
Lipidomic extracts. (**A**) Principal Component Analysis (PCA) of the lipidomic data from 5 Matyash and 5 SiMPL-DREx-extracted samples. (**B**) Heatmap of all samples, clustering based on the Ward.D2 method. (**C**) Volcano plot showing the differential expression between SiMPL-DREx and Matyash methods. Lipids with an adjusted p-value < 0.05 & a log2FC > 0.5 are colored red, those with a log2FC < -0.5 are colored blue. (**D**) Coefficient of variation of both conditions. The vertical line indicates the median CV (**E**) The ^1^H NMR spectroscopy results of Cholesterol, PC, and total fatty acid methylene –(CH2)n resonances showing about a 60:30:10 ratio extracted lipids. (**F**) Volcano plots of the no-TIC calibrated, or absolute levels, of (M1 + M2 + M3) to determine the lipid profiles of the sequential extracts results in significantly different lipid profiles for each added extraction step similar to the graph in (**E**).

To preserve RIN and limit time with RNase, the SiMPL-DREx extraction time and/or the cell lysis time is 6-fold shorter than the original Matyash^4^ and previous SiMPLEx procedures^5–9,12^. To determine potential losses due to these differences, three sequential extractions were performed with the Matyash gold standard (M1-3). **Figure 5E** are the ^1^H NMR results of the percent extraction of three representative lipids^17^, phosphatidylcholine (PC), cholesterol, and the methylene group of fatty acids (-CH_2_)_n_, in each sequential extract (M1-3). **Figure 5F** presents similar results of the deuterated lipid standards obtained by LC-MS without TIC normalization^18^. As expected, in both the ^1^H NMR and LC-MS analyses there are fewer lipids remaining with each sequential extraction. There is about 20% with each sequential extraction that would result in an 80:16:4.2 extraction ratio profile for the three sequential extracts, M1-3, but the experimental extraction profile is closer to 70:20:8 (**Fig. 5E** and **F**). This difference could be due to incomplete partitioning or equilibration between the two phases. Phosphatidylinositol is about the same in each sequential extract (**Fig. 5F**) and could suggest poor partitioning from the aqueous to the organic phase. To determine if partial extraction affects the lipid class distribution in TIC-normalized mass spectra of sequential extracts, volcano plots (**Supplementary Fig. 3B**) and lipid class comparisons between sequential extracts, M1, M2, and M3 were compared (**Supplementary Fig. 3C**). The same phospholipids, but including phosphatidylinositol, found to have relatively poor extraction efficiency by Matyash and others^10^ showed similar patterns when relative lipid abundances are compared (**Supplementary Fig. 3C**).

## DISCUSSION

The workflow in **Figure 1** shows the sequential extraction process of SiMPL-DREx for the post-mortem human brain biospecimens. The correlation of determination (R^2^) between the -omic analyses from SiMPL-DREx versus the gold standard is greater than 0.96 for all four -omics, demonstrating that SiMPL-DREx at a preliminary assessment has an extraction efficiency as good as validated single-omic extraction methods. In fact, a previous publication on tri-omic SiMPLEx methods report similar R^2^ for the metabolome, lipidome, and proteome with greater than 0.97 when compared to the similar gold standards.^5^ In this study, the transcriptome had a correlation of determination of 0.99 (**Fig. 1**) and no transcripts with significant two-fold differences between the two methods as shown in the volcano plot (**Fig. 2D**). There was no detectable RNA or DNA in upstream metabolomic and lipidomic extracts for SiMPL-DREx resulting in a pellet containing RNA that is very similar to that obtained from single-extraction of pure tissue. Although minimal, there is some RNA and DNA detected in the 80% MeOH metabolomic gold standard extraction method (**Supplementary Table 1D**). **Figure 2B** shows a PCA plot where samples cluster by donor rather than extraction method, indicating that the transcriptomic profiles are not unaffected by the upstream SiMPL-DREx lipidomic and metabolomic extraction compared with the single-step Zymo-Quick extraction. This finding is supported by the volcano plot in **Figure 2C**, in which no transcript show a significant two-fold difference between methods. Principal component 1 (PC1), which explains 82.6% of the variance, separates donors 1 and 4 from donors 2, 3, and 5, consistent with their higher RIN values (**Fig. 2B**). Previous studies have reported selective, time-dependent changes in activity and cell-specific gene expression in the human postmortem brain, which correlate with the post-mortem interval (PMI), defined as the time to brain resection and cold storage^19,20^. The separation here does not correlate with PMI as donors 1 and 4 have PMIs >22 hours and RIN values twice those of donors 3 and 5 with PMIs <12 hours (**Supplementary Table 1A** and **Fig. 2A**). The eight additional donors did not have sufficiently different PMI values to make this comparison (**Supplementary Table 1A**). Also, the correlation of clustering to RIN is confounded by sex; donors with lower RIN are female (donors 2, 3, 5) and those with higher RIN are male (donors 1, 4) (**Fig. 2A**). Recent studies have shown significant transcriptomic differences between males and females in all regions of the brain including the OP of prenatal and adult brain^21^ and this extends to the single cell level.^22^ A larger sample size containing both sexes distributed across the range of RIN values is required to unequivocally determine what factors are responsible for the two transcriptome-derived clusters.

The NIH NeuroBioBank (NBB), like many brain tissue repositories, uses RIN obtained from the occipital pole (OP) as a primary QC measure for frozen tissue samples.^23,24^ One of the goals for this SiMPL-DREx study is to maintain RIN so other -omic quality factors (i.e., DIN and PIN) and respective -omic profiles can be compared without changes caused by the extraction process. To determine if SiMPL-DREx causes adverse effects, each of the -omic extracts is compared to the respective gold standard, the single-omic extraction of tissue. The degraded RIN values in previous Folch-based^3,6,7^ biphasic multi-omic methods could be the result of exposure time between RNA and RNase in the aqueous phase during multiple upstream cell lysis and reextraction steps and extraction times. Similar to previous SiMPLEx studies SiMPL-DREx limits the aqueous exposure time to 10 min during centrifugation (**Supplementary Fig. 2**) but replaces 25 min freeze-thaw cycle in MeOH^5^ with 1 min milling time and reduces the 60 min lipid extraction time via sonication and/or orbital^5–9,12^ shaker with 10 min of vortexing. In addition, after the centrifugation step, the time in the 35% MeOH aqueous phase (**Table 1**) is minimized by isolating of the top organic phase in 45s/samples and resuspending in 80% MeOH. Resuspension in 80% MeOH is in the original SiMPLEx protocol for extraction of cells^5^, but recent penta-omic SiMPLEx protocols acquires the metabolome in the 35% MeOH aqueous fraction without addition of 80% MeOH.^6–9,12^ This is the first report of the solvent ratios of the two phases. The relative amount of MeOH remains the same at all ratios tested, but the proportion of MTBE increases in the aqueous fraction as the proportion of water deceases suggesting that MeOH is co-solvating MTBE into solution (**Supplementary Fig. 5B**). The amount of aqueous phase decreases with decreasing MTBE:MeOH:H_2_O initial extraction ratio from about 30% with 3:1:0.75 to 15% with 2:1:0.75 of total solvent volume while at 1:1:1 ratio there is no visible separation (**Table 1**). RNA and DNA are insoluble in 80% MeOH, whereas 40% MeOH will retain more nucleic acids, particularly small RNA and DNA fragments.^25^ Use of 80% MeOH in single-cell preparations has been shown to preserve RIN by precipitating both nucleic acids and RNases^26^. However, we found 6% of proteins remain in the SiMPL-DREx aqueous fraction after addition of 80% MeOH and nearly 20% of proteins remain in the 80% MeOH whole-tissue extract (**Fig. 3E**). We propose that minimizing RNA-RNase resident time during extraction and maintaining a low temperature (i.e., 4° C) is the reason for similar RIN values for each donor between the two methods (**Fig. 2A**; **Supplementary Table 1B**). Although, most of these RIN values were less than 6, a DV200 >30 is typically required for short-read RNA-seq analysis. All samples exceeded this threshold, with DV200 values above 41—even for the sample with the lowest RIN of 1 (**Supplementary Table 1C**).

The only penta-omic study examining RNA and DNA in tissues, like this study, employed a Qiagen AllPrep DNA/RNA/Protein kit.^12^ This extraction required ∼30 min of sonication and shaking in 8M urea with RNAlater followed by a ∼2.5 hr TRIzol isolation of RNA and ∼2.5 hr DNA extraction from the pellet^12^, resulting in a substantially longer workflow. In contrast, this study involves 60 s milling in Shields solution, followed by around 60 min of total RNA and DNA extraction using Zymo Quick column-based RNA protocols (**Supplementary Fig. 2** and **Fig. 5**) reducing DNA and RNA isolation by an hour. In the present and previous SiMPL-Ex studies^6,12^, tissue processing involved pulverization with a mortar and pestle under liquid nitrogen, a widely accepted standard for RNA extraction. However, recent work indicates that this method can produce variable RIN values compared to bead milling or mechanical homogenization.^27^ In short, based on our results (**Fig 2A, Supplementary Table 1B**) and others^12^, tissue homogenization is likely the primary cause of the decrease in RIN prior to the extraction procedure.

In contrast to the RIN results, the DIN values were slightly degraded (**Fig. 2A–** DIN table). Sonication and vortexing results in mechanical shearing of DNA.^28^ The Zymo-Quick protocol calls for a shorter sample vortex period (∼1 min) compared to SiMPL-DREx (20 min), and it is likely that the added sample vortex time increased shearing of DNA, resulting in poorer DIN values. In fact, the pellet from Donor 3, which has the biggest difference in DIN from whole tissue (**Fig. 2A–**DIN table) had the longest vortex time because the pellet would not dissociate without the help of a probe to break it up and extensive trituration. Overall, these results validate the SiMPL-DREx method to extract RIN invariant transcriptome from the pellet with solid-phase extraction, Zymo-Quick, as compared to single-omic extraction of tissue. The slight degradation of DNA from the SiMPL-DREx extraction may not negatively impact many applications, such as short-read high throughput sequencing. However, one may need to consider single-omic extraction of DNA directly from tissue to maximize DNA length if technologies requiring high molecular weight DNA (such as long-read sequencing) are to be used.

The proteomic results are shown in **Figure 3**. Similar to the DNA and RNA extraction, protein extraction was performed on human brain tissue subjected to either the SiMPL-DREx workflow or a standard single-omic whole tissue control. In contrast to the RIN and DIN trends, the Protein Integrity Number (PIN) exhibited no variance between donors 1 and 4 and donors 2, 3, and 5 (**Fig. 3A**) suggesting uniform degrees of protein degradation across all samples. Principal component analysis (PCA) of the proteomic data revealed slight clustering based on the extraction method rather than by donor (**Fig. 3B**) - a trend unique to this omic layer. This is likely due to the 6% protein loss in the SiMPL-DREx pellet compared to the single-omics whole-tissue method (**Fig. 3E**) since the coefficient of variation was marginally lower for the SiMPL-DREx cohort (**Fig. 3D**), and the volcano plot showed no proteins with >2 fold significant change (adjusted *p*-value < 0.05) between methods (**Fig. 3D**). Overall, there was no significant difference between the proteome obtained from the SiMPL-DREx pellet and tissue (**Fig. 3D**).

Conversely, the gold standard 80% MeOH metabolomic extraction resulted in a ∼20% proteins loss. While 80% MeOH is an established protein precipitation and metabolomic solvent it has previously been found to contain enzymatically active proteins.^29^ Alternative mixtures (e.g., MeOH:acetonitrile:water) have recently demonstrated superior metabolomic performance.^30^ Unexpectedly, although the SiMPL-DREx metabolomics phase similarly utilizes 80% MeOH, it exhibited a four-fold reduction in protein loss compared to direct 80% MeOH tissue extraction (**Fig. 3E**). This enhanced retention is likely driven by the initial SiMPL-DREx step; treating the tissue with 3:1 MTBE:MeOH may denature the proteins and invert hydrophobic residues, causing irreversible precipitation during the subsequent metabolomic extraction phase.^31,32^ In addition, there is about 20% MTBE in the aqueous fraction prior to MeOH addition that could aid protein precipitation especially when 80% MeOH is added to the aqueous phase prior to extraction (**Fig. 6**).

The metabolomic results show little separation between the methods in the PCA plots **(Fig. 4A**) but rather separates by donor like the transcriptome PCA plot (**Fig. 2B**), with donors 1 and 4 separated by PC1 from donors 2, 3, and 5 and separated by cluster analysis (**Fig. 4B**). Donors 1 and 4 cluster and have higher RIN but are confounded by sex (**Fig. 2A**). In fact, metabolomic differences between sexes have been found in the brains of Alzheimer’s disease mouse models^33^. Several metabolites have been correlated with post-mortem times. Essential amino acids, like tryptophan have been proposed to be a quality factor^34^ and used as measures of UPR and autophagy^35^ because they cannot be synthesized and derive from protein degradation. A more in-depth study of RIN and metabolomic tissue quality markers is required to more clearly understand factors driving the clustering observed in the multiple -omic datasets.

The volcano plot (**Fig. 4C**) shows some metabolites differ between methods. In addition, **Figure 4D** shows a slight leftward shift in the %CV distribution for SiMPL-DREx, indicating marginally improved reproducibility. The primary distinction between the biphasic SiMPL-DREx and the monophasic 80% MeOH gold standard methods is that a greater fraction of nonpolar metabolites can partition into and be lost within the organic phase (**Fig. 4E**).^5,9^ This loss correlates with hydrophobicity of the amino acids^5^ and can be predicted for physicochemical properties like the octanol–water partition coefficient (LogP) and the Hansch hydrophobic parameter (π) shown in **Figure 4F**. The extent of polar metabolite loss is also influenced by the solvent-to-tissue ratio^36^, MTBE:MeOH:H_2_O solvent ratios^7,9^, and use of ionic salts/pH^36,9^. A higher solvent-to-tissue ratios can improve both extraction efficiency and reproducibility for abundant lipids such as phosphatidylcholine (PC), with a 100:1 ratio outperforming a 20:1 ratio in human plasma.^37^ A lower solvent-tissue ratio would decrease polar metabolites at the expense of lower lipid extraction efficiencies. A ratio of 50 µL solvent per 1 mg wet human brain tissue was used in this study which is similar to solvent-tissue ratios of previous SiMPLEx studies extracting solid tissues.^6–9^ Hemmati and others^9^ compared MTBE:MeOH:H_2_O solvent ratios 10:3:2.5 and 7:3:2.5 and found the former to have a slightly higher recovery for both polar and nonpolar metabolites, but the mean recovery efficiencies of all classes of polar metabolites were not significantly different (*p*<0.05). Sostare and others^7^ found a ratio of 2.6:2.0:2.4 to perform best for extracting the polar metabolites. A ratio of 8.5:2.83:2.5 was used in this study - a ratio between the two tested by Hemmati and others.^9^ Decreasing pH or adding small amounts of ion-pairing salts can increase lipid extraction efficiency; however, these conditions may also enhance the loss of hydrophobic metabolites from the aqueous fraction.^36^ Ion-pair extraction—defined as the transfer of ionized solutes into an organic phase through the addition of an oppositely charged ion—has been widely used for decades.^36^ The original protocol reported by Matyash and others^4^ and Coman and others^5^ included 0.1% ammonium acetate (13 mM), during tissue homogenization or phase separation, respectively, which functions as an ion pairing organic salt. A recent comprehensive evaluation of multiple lipid extraction methods across various tissues compared the Matyash protocol with and without 10 mM ammonium acetate and demonstrated that the ion-pairing reagent significantly increased both the number and abundance of detected lipids, but the effect on polar metabolites like amino acids was not measured.^38^ Zwitterionic species can form ion pairs with ammonium acetate, thereby increasing their solubility and thereby loss to the organic phase. Using much higher concentrations of an inorganic salt, 300 mM ammonium chloride, Hemmati and others^9^ were able to salt out lipids and salt in polar metabolites into the organic and aqueous phases, respectively. For this study, we did not want to taint the future metabolomic analysis of acetate by inclusion of ammonium acetate and avoided high inorganic salt loads to avoid possible LC-MS ionization issues. However, future studies may consider including these options to increase extraction efficiencies.

Since SiMPL-DREx starts with a biphasic Matyash separation one would expect identical results, and any difference would likely reflect technical and experimental variations. Although the donors 1-5 cluster in the PCA plots (**Fig. 5A**), the clustered heatmaps (**Fig. 5B**) show high dissimilarity of donor 3 and less so of donor 2 by method which correlated better with PC3 and PC4, respectively (**Supplementary Fig. 3A**). However, volcano plot (**fig. 5C**) shows no lipids out of 391 that were significantly higher in the SiMPL-DREx method. The % CV plot in **Figure 5D** shows the SiMPL-DREx method with higher average %CV’s with an upward shift in profile. In comparison to the lipidome of laboratory rodent brain, the larger average %CV of the post-mortem human brain could be a real effect caused by both genetics and environmental divergence, and especially due to post-mortem effects on the lipidome.^39^

To determine the residual lipids left after the first extraction on the absolute and TIC-normalized lipid profiles were determine in the sequential extracts (M1-3) shown in **Figure 5E** and **5F** and **Supplementary Figure 4C**, respectively. The extraction efficiencies were similar for the sequential with the deuterated standard except for phosphatidyl inositol and similar results were found here (**Fig. 5E** and **5F**). Overall, subsequent extractions had similar TIC-normalized lipid profiles except for those lipids with lower concentrations (**Supplementary Fig. 4C**). This difference could be due to the more concentrated lipids contributing less overall to the total ion current (TIC) thereby over-estimating the lower concetration lipids in subsequent extracts, or the one sixth shorter extraction time during the initial MTBE:MeOH lipid extraction step did not permit sufficient dissolution.

In conclusion, shortening the cell disruption and lipid extraction time while maintaining low temperature (4° C) maintains RIN in post-mortem human brain. There was no significant difference on transcriptome, proteome, or lipidome profiles between SiMPL-DREx and the gold standard methods. The SiMPL-DREx 80% MeOH metabolomic fraction had one fourth the protein loss compared to the gold standard, 80% MeOH. Despite a similar 80% MeOH extraction solvent, the superior protein extraction efficiency by SiMPL-DREx may be due to protein denaturation during the upstream 10 min MTBE:MeOH vortexing step and subsequent centrifugation to pellet proteins. The metabolome had some metabolite differences from the 80% MeOH gold standard metabolomics extraction. Disproportionate loss of more hydrophobic amino acids partitioning into the upper organic layer was shown to correlate with the compound’s extraction coefficient (K_ex_) and could be responsible. This is the first time the solvent ratios have been quantified in the organic and aqueous fractions using ^1^H NMR spectroscopy. The proportion of MTBE in the aqueous fractions increases with increasing MeOH proportion and decreasing water proportion likely caused by co-solvation with MeOH. The interplay of extraction conditions that help one omic will affect another. In this study, the SiMPL-DREx protocol with 50:1 solvent-to-PMI human brain weight ratio and a 11-minute cell disruption and lipid extraction time is an efficient and valid method for extracting most lipids while minimizing RNA, DNA, metabolome and protein losses and maintaining RIN in a single biospecimen.

## ON-LINE METHODS

### Materials

Methanol ACS grade (>99.8% pure), RNase-free water, methyl-tertbutyl ether, deuterated acetonitrile (98.8% pure), deuterium oxide (98.8% pure), and 5mm NMR tubes were purchased from MilliporeSigma. Qubit RNA assay kit was purchased from ThermoFisher Scientific Inc., RNA screen tape and buffer was purchased from Agilent Technologies Inc. Quick-DNA/RNA MiniPrep Plus kit and 2 mm milling tubes with beads was purchased from Zymo Research Corporation.

### Selection of Human Brain Samples

Post-mortem human brain tissue from the occipital pole were obtained from the NIH NBB. Upon dissection, samples were frozen either in liquid nitrogen or a slurry of isopentane and dry ice and stored at -80° C. Five neurologically unaffected control donors proportional to the population of three males and two premenopausal women with ages less than 55 years. Eight additional neurologically unaffected control donors were subsequently obtained for additional RIN and DV200 measurements. Given the measurement of metabolic, lipid, and proteomic differences, individuals with metabolic diseases that impact the nervous system such as diabetes and liver disease, autoimmune neurological conditions such as MS or lupus, and systemic infection with hepatitis B, C, or HIV were excluded. To obtain the most rapid antemortem death of agonal duration less than 1 hr the manner of death was natural and the cause of death was most commonly cardiorespiratory arrest. The metadata for each of the thirteen donors are given in **Figure 1A**.

### Optimizing Extraction Conditions

The effect of extraction solvent composition and sample mass on the final MTBE:MeOH:H_2_O ratios of the organic and aqueous phases has not previously been characterized for biphasic multi-omic extractions, despite its importance for determining extraction selectivity. Five samples were made in MTBE:MeOH:H_2_O ratios of: (a) 900μL:300μl:250μl (62:21:17), (b) 750μL:250μl:250μl (60:20:20), (c) 700μL:300μl:250μl (56:24:20), (d) 667μL:333μl:250μl (57:35:18), (e) 330μL:330μl:330μl (33:33:33). Solvents and samples were on ice, and once mixed they were vortexed for 20 s, and centrifuged at 15,000g for 10 min at 4° C. Samples of 6 μL of organic and aqueous phases were added to 594 μl of deuterated acetonitrile (d_3_-ACN), transferred to 5 mm NMR tubes and analyzed on a Bruker Avance III HD 700 MHz four channel spectrometer equipped with a TCI H-C/N-D 5 mm CryoProbe at 298° K. A single pulse sequence (Bruker zg) with no solvent suppression, digital quadrature detection (DQD), and an acquisition time of 2.93 s plus interpulse delay of 30 s resulting in a 32.93 s repetition time with 8 transients. The sweep width was 11,160 Hz generating 65,536 complex points FID resulting in a 32,768 real spectrum after Fourier transformation. TopSpin v.3.6.3 was used to process and analyze spectra. The FID was zero filled to131,072 points with a line broadening of 0.8 Hz exponential and 2 Hz shifted sine bell, phased and base line corrected the same. The peaks were integrated and normalized to the residual acetonitrile set to 1.0. Representative ^1^H NMR spectra of d-ACN (bottom), aqueous (middle) and organic (top) integrations are shown in **Supplementary Fig. 5A** and generated the MTBE:MeOH:H_2_O ratios shown in the **Table 1**. All solvent were within several-fold of each other so single peak experience radiation dampening. Water is about 5% soluble in MTBE and vice versa. To determine whether there is a correlation between higher methanol content and increased co-solvation of MTBE in the aqueous phases, the ratio of MTBE and MeOH volume fractions in the aqueous phase (MTBE/MeOH % in Aq.) is plotted against the total water fraction (%) in the aqueous phase. There is a linear correlation that reflects not only MTBE solubility in water, but the same percentage of MeOH (∼35%) competing between H-bonding with water versus dissolving MTBE (**Supplementary Fig. 5B**). This result is reflected in the 2:1:1 ratio generating a 2-fold smaller aqueous phase out of the total volume compared to the higher 3:1:0.75 ratio and and the 1:1:1 ratio had no detectable phase separation. To determine possible mixing of the two phases at the interphase, sequentially, the top and bottom halves were transferred, then the remaining residual aqueous was transferred including part of the interphase, and then the remaining organic layers was transferred (**Supplementary Fig. 2C**). Only the aqueous phase that contained part of the interphase differed from the bottom half of the aqueous phase. Therefore, in an effort to minimize RNA-RNase residence time in the aqueous fraction and maintain and organic transfer time of <45s/sample, the final workflow left ∼150 μL of the upper organic layer prior to the addition of 80% MeOH to the aqueous fraction for the metabolomic extraction (**Supplementary Fig. 2**).

### SiMPL-DREx and Gold Standard Extraction Workflows

For all extractions, the thirteen donor brain samples were powderized using a mortar and pestle with constant liquid nitrogen addition, stored at -80° C until used when 25 mg was weighed for each of the following extraction workflows.

#### Solvent Volume-to-Tissue Ratios

The solvent volume (μL)-to-wet tissue weight (mg) was 2.5-fold higher (54μL:1mg wet human brain) than Matyash^4^ (**Supplementary, Fig. 2** and **Fig. 4**) but similar to previous SiMPLEx studies using tissues. ^5–9,12^ The SiMPL-DREx solvent-to-tissue ratio for the second step to extract the metabolome was 50 μL:1mg, similar to the 60:1 ratio used in 80% MeOH metabolomic gold standard extraction method using mouse brain^40^. For RNA and DNA extraction using Zymo-Quick columns 4.2 ± 1.2 mg was taken from the SiMPL-DREx pellet leaving 13.4±3.5 mg for proteomic analysis.

#### SiMPL-DREx Extraction

The detailed SiMPL-DREx extraction workflow is provided in **Supplementary Fig. 2**. The primary objective of the extraction is to maximize RNA integrity number (RIN) by minimizing RNA exposure to RNases. This is achieved through a rapid, single-step lipid extraction performed under cold conditions, with addition of MTBE and methanol (MeOH) stored at −20 °C, 1 min of cell milling, and transfer to a microfuge tube. About 50 μL remains in the microfuge tube depending on the number of beads if not collected with a short quick low speed centrifuge step. This loss affected the final ratio of MTBE:MeOH:H_2_O from 9:3:2.5 to 8.5:2.83:2.5. The samples were vortex at 3200 rpm at 4 °C for 10 min, then samples were placed on ice and 250 μL of DEPC water to induce phase separation (**Supplementary Fig. 2**). After vortexing, the samples were centrifuged at 15,000g and 4 °C to precipitate RNases and other proteins away from the aqueous phase to the pellet. Previous SiMPLEx methods used a range of centrifugation speeds from 2,000-20,000 g^5–9,12^ for 5-10 min but Qiagen protocol to isolate RNA [https://www.thermofisher.com/us/en/home/life-science/dna-rna-purification-analysis/rna-extraction/rna-sample-extraction.html?open=tissue#tissue] gives a range of 12,000-16,000 g, so the upper limit was chosen to further increase RNase precipitation (15,000g for 10 min). To insure processing was less than 45s/sample, 850 μL of top organic layer was rapidly transferred to a microfuge tube leaving about 150 μL of top and interphase and this was later partitioned into 4 different microfuge tubes of varying amounts for either NMR metabolite and lipid analysis, LC-MS lipidomic analysis, RNA/DNA and proteins quantification (**Supplementary Fig. 2**). After about 4 min for the 5 initial samples and and 6 min for the 8 additional samples plus blanks for transfer of the upper organic fraction, 800 μL MeOH is added to the ∼500 μL of aqueous phase plus interphase fraction, resuspended, then centrifuged at 15,000g for 10 min. The supernatant of about 1.2 mL is transferred to four different microfuge tubes for the same quantification as described above (**Supplementary Fig. 2**). About a fourth of the pellet was taken for the RNA and DNA analysis using the protocol described below for Zymo-Quick solid phase extraction, while the remainder of the pellet was used for protein analysis.

#### Gold Standard Lipidomic Extraction from Tissue

The same SiMPL-DREx workflow, shown in **Supplementary Fig. 3**, starts the same as the gold standard Matyash extraction^4^ workflow detailed in **Supplementary Fig. 4**, but the Matyash first extraction step is followed by two sequential lipid extractions while leaving the residual aqueous phase from the first step (**Supplementary Fig. 4** - left flow chart). In the second extraction a total of 965 µL MTBE:MeOH:H_2_O (9:3:2.5) was added, resuspended with vortexing and titurating, and then the top organic layer was portioned into 4 different microfuge tubes of varying amounts for either NMR metabolite and lipid analysis, LC-MS lipidomic analysis, and RNA/DNA and protein quantification (**Supplementary Fig. 4**). In the third extraction, because there was about 500 µL of residual aqueous phase, the solution was washed with 670 µL MTBE:MeOH (3:1), vortexed for 1 min, and a smaller amount was portioned into 4 microfuge tubes (**Supplementary Fig. 4**).

#### Gold Standard Metabolomic omic Extraction from Tissue

The detailed gold standard 80% MeOH extraction workflow is provided in **Supplementary Fig. 4** (right flow chart). There were three sequential extractions with volumes shown in **Supplementary Fig. 4** and the pellet was again resuspended by vortexing titurating at each step. Then the supernatant was partitioned into 4 different microfuge tubes of varying amounts for either NMR metabolite and lipid analysis, LC-MS lipidomic analysis, and RNA/DNA and protein quantification (**Supplementary Fig. 4**).

#### RNA/DNA Zymo-Quick extraction from Tissue

Total RNA and genomic DNA were isolated from brain tissue using the Zymo Research Quick-DNA/RNA extraction workflow (e.g., Quick-DNA/RNA Miniprep Plus–type chemistry), following the manufacturer’s protocol with minor modifications to optimize performance for lipid-rich neural tissue. Briefly, frozen brain tissue was maintained on dry ice and a defined mass of tissue (typically 10–30 mg) was transferred to pre-chilled tubes for rapid homogenization in the kit-provided lysis buffer. Tissue disruption was performed by mechanical homogenization (e.g., bead-based homogenizer or rotor–stator) for 60 s. Lysates were clarified by brief centrifugation to reduce insoluble debris, and the supernatant was processed through the column-based workflow to enable sequential purification of RNA and DNA from the same sample input. RNA was purified using silica-membrane binding and washed to remove proteins, lipids, and inhibitors; on-column DNase treatment was applied when specified by the kit protocol to minimize genomic DNA carryover into RNA preparations. Genomic DNA was purified from the corresponding fraction using the kit’s DNA-binding conditions and column cleanup, yielding high-molecular-weight DNA suitable for downstream library preparation.

Purified nucleic acids were eluted in nuclease-free water (or kit elution buffer) and stored at −80 °C (RNA) and −20 °C (DNA) until use. RNA concentration was measured fluorometrically (Qubit RNA BR), and RNA integrity was assessed by capillary electrophoresis (TapeStation) using RIN as the primary metric in addition to DV200. DNA concentration was measured fluorometrically (Qubit dsDNA BR), and DNA integrity was evaluated by electrophoretic profiling (TapeStation Genomic DNA) and/or fragment analysis of a long amplicon when needed. Negative extraction controls were included per batch to monitor reagent/background contamination.

#### Proteomics Sample Preparation

Each tissue was lysed in 15 μL per mg of 8M urea, 50mM Tris pH 8, 1% SDS, supplemented with 1x protease inhibitors. Pre-chilled beads were added to each sample, then subject to bead-beating at speed 3 in 30-second pulses for 5 minutes. Lysates were centrifuged at 15,000 xg for 15 minutes at 4 °C, and supernatants were transferred to clean tubes. Proteins were incubated with 6 volumes of ice-cold acetone overnight at -20 °C. The following day, samples were centrifuged at 15,000 xg for 15 minutes at 4 °C, supernatant was removed, and the pellet was washed with 100 μl of ice-cold acetone. Pellets were air dried at room temperature for 10 minutes before being resuspended in 500 μl of AccelerOme lysis buffer (Thermo Scientific). Protein concentration was determined using a BCA assay. For each sample, 70 μg of protein was loaded onto an AccelerOme plate; if there was less than 70 μg of protein for a sample, then the entire sample was used. Samples were further prepared using the Thermo Accelerome automated platform with the label-free Accelerome preparation kit. The workflow included reduction with DTT, alkylation with iodoacetamide, LysC/trypsin digest at 37 °C, desalting, and absorbance-based peptide quantification. Peptides were dried via vacuum centrifugation, then stored at -80 °C until further analysis.

### -Omics Data Analysis

For the transcriptomics, proteomics, metabolomics, and lipidomics data, each was filtered for having at least 3 valid values in one condition. Figures were generated in R (4.5.1) using the ggplot2 package. Date was analyzed by paired t-test between extraction methods of the same donors using FDR, or the q-value.^41,42^ Adjusted p-values were calculated using the Benjamini-Hochberg method to correct for multiple comparisons. Analytes with a q-value < 0.05 were considered statistically significant.

### Transcriptomics and Data Analysis

Transcriptomics library preparation and sequencing: Total RNA extracted from brain tissue was used to generate bulk RNA-seq libraries using the Watchmaker Genomics RNA-seq library preparation workflow with Polaris rRNA depletion to remove ribosomal RNA and enrich for informative transcript species. Libraries were quantified fluorometrically and assessed for fragment size distribution by capillary electrophoresis prior to pooling. Pooled libraries were sequenced on a DNBSEQ-T7 instrument to generate paired-end 2×150 bp reads (FASTQ format) for downstream analysis.

RNA-seq processing, alignment, and differential expression analysis (CLC Genomics Workbench v25): All computational analyses were performed using CLC Genomics Workbench v25 (QIAGEN). Raw FASTQ files were imported into CLC and processed using the built-in read trimming tool. Trimming parameters were: quality-score trimming enabled (quality limit = 0.05); automatic read-through adapter trimming enabled (custom “New trim adapter list”); trimming of ambiguous nucleotides disabled; 3′ homopolymer trimming enabled for polyA only (polyA = Yes; polyC/polyG/polyT = No); no removal of fixed numbers of 5′ or 3′ terminal nucleotides; and no trimming to a fixed length (maximum length = 150; trimming from the 3′ end). Trimming was applied to both reads in a pair (first read = Yes; second read = Yes). Reads were not discarded based on length, and reports were generated.

Trimmed reads were mapped using the CLC RNA-Seq Analysis pipeline to an annotated genome reference (hg38, reference type: genome annotated with genes and transcripts; gene track: *hg38_Gene*; mRNA track: *hg38 (RNA)*). Alignment settings were: mismatch cost 2, insertion cost 3, deletion cost 3, length fraction 0.8, similarity fraction 0.8, global alignment disabled, and maximum number of hits per read 10. Libraries were analyzed as strand-specific (reverse) bulk RNA-seq. Broken pairs were ignored, paired reads were not counted as two, and spike-in controls were not used. Gene-level expression was quantified as total counts, and normalized expression values were reported as TPM (transcripts per million). A mapped reads track and analysis reports were generated; lists of unmapped reads were not generated.

### Proteomics and Data Analysis

#### Proteomics LC-MS/MS Analysis

All samples were reconstituted in 2% acetonitrile with 0.1% formic acid and normalized to 0.1 µg/µl. 2 μL were taken from each sample and combined to create a pool sample that was run before and after the sample sequence assess technical reproducibility. Samples were analyzed via LC-MS/MS using a Vanquish Neo coupled to an Orbitrap Astral mass spectrometer (Thermo Scientific). Samples were injected onto an Aurora® Ultimate TS column (75 μm id × 25 cm, 1.7 μm particle size; IonOpticks) and separated over a 30 min gradient. The separation gradient consisted of 5-45% mobile phase B at a 300 nl/min flow rate, where mobile phase A was 0.1% formic acid in water and mobile phase B was 0.1% formic acid in 80% acetonitrile. MS1 full scans (m/z 380-980) were acquired with a resolution of 240,000 in the Orbitrap analyzer with a maximum injection time of 5 ms and an AGC target of 500%. MS2 scans were acquired using the Astral detector operated in Data-Independent Acquisition (DIA) mode, covering a scan range of m/z 150-2000 with a cycle time of 0.6 s. Isolation windows were set to 3 m/z with a higher collision dissociation (HCD) energy of 25%, an AGC target of 500%, and a maximum injection time of 3 ms.

#### Proteomics LC-MS/MS Data Analysis

Raw data files were processed using Spectronaut (v20; Biognosys) and searched against the Uniprot reviewed human database (containing 20,434 entries, downloaded January 2025), appended to the MaxQuant contaminants database (containing 246 entries). The following settings were used: enzyme specificity set to trypsin, up to two missed cleavages allowed, cysteine carbamidomethylation set as a fixed modification, methionine oxidation and N-terminal acetylation set as variable modifications. A false discovery rate (FDR) of 1% was used to filter all data. Imputation was disabled, and single hit proteins were retained. Paired student’s t-tests were conducted, and p-values, FDR-corrected p-values (q-values), along with log2 fold change ratios were calculated in Spectronaut. Proteins with an absolute log2 fold change ≥ 0.5 and a q-value < 0.05 were considered significant.

#### Proteomic Analysis of Molecular Weight, Hydrophobicity, and Isoelectric point

The molecular weight (MW), isoelectric point (pI), and hydrophobicity of identified proteins were calculated in R (v4.5.1) using the Peptides package (v2.4.6).^43^ Protein sequences were extracted from the same FASTA database used for proteomics database searching. Hydrophobicity was calculated using the Kyte-Doolittle scale. All plots were generated using the ggplot2 package (v4.0.0) in R.

#### BCA Analysis of Protein Concentration

Aliquots from the various extracts from the five donors (n=5) were obtained for BCA analysis. Samples were dried by Speed-Vac and stored at -80° C until analyzed. All samples were resuspended in 50 μL AccelerOme Lysis buffer (Thermo), protein quantitation was conducted using a Colorimetric BCA assay (Pierce) and analyzed using Excel. A paired t test was performed between SiMPL-DREx and gold standard samples and the results are shown in **Figure 3E**.

### PIN Calculation

The PIN calculation code was written in R with help from ChatGPT using the algorithm and formulas published in the Shao W et.al publication.^13^ The raw data were searched in Spectronaut (v20; Biognosys) using the same settings as described above, except the enzyme specificity was set to semi-specific trypsin. For each protein within each replicate, we calculated an intensity-based Proteome Integrity Score (iPIS) by summing the total peptide ion intensity and the intensity attributable specifically to canonical tryptic peptides. The iPIS was defined as 1 − (tryptic peptide intensity/total peptide intensity), such that higher values reflect a greater proportion of non-tryptic peptides. The Proteome Integrity Number (PIN) for each replicate was then computed as the mean iPIS across all quantified proteins.

To assess whether any samples exhibited abnormal degradation, the distribution of PIN values was modeled using a Weibull distribution. Weibull shape and scale parameters were estimated by maximum likelihood, and replicate-level p-values were calculated from the model’s cumulative distribution. An iterative procedure was used to refine the null distribution: samples with p < 0.02 were temporarily excluded, the Weibull distribution was refit, and p-values were recalculated until no further samples fell below the threshold. The resulting p-values and inclusion status were used to identify replicates consistent with non-degraded proteomes.

### Metabolomics (LC-MS and NMR) and Data Analyses

#### Metabolomic LC-MS/MS Analysis

Samples were analyzed using a Q Exactive HF-X (ThermoFisher, Bremen, Germany) mass spectrometer coupled with a Waters Acquity UPLC liquid chromatograph system. Samples were introduced via a heated electrospray ionization (HESI) source using polarity switching at a flow rate of 0.25 mL/min. Electrospray source parameters were set as follows: spray voltage 3.5 kV, sheath gas (nitrogen) 53 arb, auxiliary gas (nitrogen) 14 arb, sweep gas (nitrogen) 2 arb, nebulizer temperature 400 °C, and capillary temperature 300 °C. Analyses were conducted from *m/z* 200-2000 in full scan mode and data dependent MS2 at a resolution setting of 60,000. Separations were conducted on a Waters Acquity UPLC BEH -HILIC column (2.1 x 100 mm, 1.7 um particle size) using conditions previously described. ^45^

#### Metabolomic LC-MS/MS Data Analysis

Data acquisition was performed using Xcalibur software (ThermoFisher, Bremen, Germany). Compound Discoverer (v 3.3 ThermoFisher, Breman, Germany) was used to identify metabolites. Method parameters were identified by accurate mass (mass error of precursor = 5 ppm). Samples were grouped in Compound Discoverer and submitted for identification using Thermo’s workflow titled “Untargeted Metabolomics with Statistics Detect Unknowns with ID using Online Databases”. The following databases were used for identification: mzCloud, ChemSpider, and Metabolika. Tissue mass was utilized for data normalization. Data visualization was completed utilizing MetaboAnalyst 6.0.

#### NMR Metabolomic Analysis

After deuterated chloroform extracted the lipid extracts for lipidomic analysis, the microfuge tubes were vacuum dried. The residue was dissolved in 600 μL of D_2_O containing 0.05 mM trimethylsilyl-2,2,3,3-tetradeuteropropionic acid (TSP) and transferred to a 5 mm NMR tube for high resolution NMR analysis. The 1D ^1^H NMR spectra were acquired at 25 °C using a 19.97 T Bruker Avance III HD 850 MHz spectrometer equipped with a TCI H-C/N-D 5 mm CryoProbe with a zgesgp pulse sequence using a 90° flip angle, with a 1.5 s excitation sculpted selective 180° shaped pulses to eliminate water, a 1 s acquisition time and 3 s relaxation time resulting in a 4 s repetition time. The sweep width was 17007 Hz and acquired with 32,000 complex points, and 2048 transients. All NMR spectra were processed using Chenomx Inc. version 12.0 software (Edmonton, Alberta). Imported FIDs were zero-filled to 256,000 points, and an exponential line broadening of 1 Hz was applied before Fourier transformation. Phase and baseline correction were conducted for the spectra. Resulting concentrations were exported to Excel for data processing.

### Lipidomics and Data Analysis

#### Lipidomic LC-MS/MS Analysis

Samples were dried by vacuum and reconstituted in 100 µL of isopropyl (IPA) alcohol for analysis. Analysis was performed using a Thermo Q Exactive HFX coupled to a Waters Acquity H-Class LC. A 100 mm x 2.1 mm, 1.7 µm Waters BEH C18 column was used for separations. The following mobile phases were used: A- 60/40 acetonitrile (ACN)/H_2_O, B- 90/10 IPA/ACN; both mobile phases had 10 mM ammonium formate and 0.1% formic acid. A flow rate of 0.2 mL/min was used. The starting composition was 40% B, which increased to 95% B at 12 min (held until 15 min) then 40% B at 15.1 and held for 5 mins for re-equilibration. Samples were analyzed in positive/negative switching ionization mode with top 5 data dependent fragmentation.

#### Lipidomic LC-MS/MS Data Analysis

Raw data was analyzed by LipidSearch 5.0 software. Lipids were identified by MS2 fragmentation (mass error of precursor = 5 ppm, mass error of product = 8 ppm). The identifications were generated individually for each sample and then aligned by grouping the samples (SiMPL-DREx vs Matayash extraction #1 vs #2 vs #3). The lipid identities were curated with a custom r script where sequence of filters are performed. First, erroneous Lipid ID duplicates with the same peak area values are removed, non-randomly missing values are set to zero, and selected adducts of cholesterols (+H-H2O), all the protonated, deprotonated, and ammoniated adducts for glycerides (+H, -H, +NH4), and only the protonated and deprotonated adducts of all other lipids (+H, -H). Then lipid identieis are filter by expected chemical trait space whereby each lipid class has an expected number of carbons, number of double bonds, and retention time. Approximately one third of lipids detected had multiple features assigned the same identification within a sample and these were filter by features of the XIC area and retention time. These multiplicands could be true signals due to bimodal peaks, or they could be false signals, baseline “wobbles” which were incorrectly identified as spectra. Usually, false signals are relatively small and have earlier or later than average retention times. For each lipid/sample, the median feature area and retention time is calculated, then iteratively removed small, distant peaks with an area smaller than the median and a retention time further from the median retention time than the median temporal difference, until no additional features meet the criteria for removal. For remaining multiplicands (usually only duplicates), only the largest peak is retained if features were far from each other, or features were summed if retention times were close. Such features were characterized as near versus far using the median difference in retention time across remaining multiplicands as the threshold. Once spectral features are filtered down to a single peak per lipid per sample, sample differences are corrected in tissue mass, cell count, and/or extraction dilution. Although class specific adducts are filtered in the first step some lipid classes with multiple potential suitable adducts may remain. For these the total spectral peak areas are calculated for each adduct and only the adduct with the largest total signal is retained. Our datasets include missing not at random values representing lipids that were not detected in some samples. To enable comparisons across samples, values were imputed using the half-minimum approach and then log2-transform all data, centered and scaled the data for each lipid in the dataset to enable comparisons across lipids. The resulting data are exported to csv files for further analysis and manipulation.

#### Lipidomic NMR Analysis

Dried samples were redissolved in 600 mL of deuterated chloroform containing 0.05% (V/V) tetramethylsilane and transferred to a 5 mm NMR tube for high resolution NMR analysis. The 1D ^1^H NMR spectra were acquired at 25 °C using a 19.97 T Bruker Avance III HD 850 MHz spectrometer equipped with a TCI H-C/N-D 5 mm CryoProbe with a zgesgp pulse sequence using a 90° flip angle, with a 1.5 s excitation sculpted selective 180° shaped pulses to eliminate water at 4.7 ppm, a 1 s acquisition time and 2 s relaxation time resulting in a 3 s repetition time. The sweep width was 17007 Hz and acquired with 32,000 complex points, and 1024 transients. All NMR spectra were processed using Bruker TopSpin version 3.6.3 software (Edmonton, Alberta). Imported FIDs were zero-filled to 131,000 points, and an exponential line broadening of 1 Hz was applied before Fourier transformation. Phase and baseline correction were conducted for the spectra. Selected peaks were integrated. Resulting peak area values were exported to Excel and calibrated to the TMS peak area to obtain molar ratios.

## Supporting information

Penta-omic Supplemental Figures

## Data Availability

The script used in this study to generate figures and calculate the PIN values is available at: 10.5281/zenodo.20765389. The proteomics dataset generated and analyzed during the current study has been deposited in the PRIDE repository (ProteomeXchange Consortium) under the accession number PXD080003. All other data supporting the findings of this study are available from the corresponding author upon request. The RNAseq data generated and analyzed for the current study has been deposited in the prior to submission. The metabolomic and lipidomic data is available at the NIH Common Fund’s National Metabolomics Data Repository (NMDR) website, the Metabolomics Workbench, https://www.metabolomicsworkbench.org prior to submission.

## ACKNOWLEDGMENTS

The work is supported by grants from NIA (R21AG088936) and the UNC School of Medicine Core Facilities Advocacy Committee program. The High Throughput Sequencing Facility service is supported in part by an NCI Center Core Support Grant (P30CA016086). This research is based in part upon work conducted using the UNC Metabolomics and Proteomics Core Facility, which is supported in part by NCI Center Core Support Grant (2P30CA016086-45) to the UNC Lineberger Comprehensive Cancer Center and Nutrition and Obesity Research Center (P30DK056350). The UNC Department of Chemistry Mass Spectrometry Core Laboratory used the Q Exactive HF-X LC-MS system for metabolomic and lipidomic analyses, which is supported by the National Science Foundation under Grant No. (CHE-1726291). Human tissue samples included in this study were provided by the University of Pittsburgh, Maryland, and Harvard Brain and Tissue Repositories of the NIH NeuroBioBank. We express our gratitude to all the brain donors and their families whose selfless generosity made this research possible. We appreciate Savannah Weaver’s help with the r script for lipidomic annotation.

## Author Contributions

PMM and JMM conceived the experiment. PMM, JMM, SPL, LEM, WKS, BME all contributed to the design of the experimental procedure. PMM performed and analyzed all transcriptomic and genomic studies. LEM, SPL, and TSW performed and analyzed all proteomic studies. BEM performed and analyzed LC-MS studies; JMM performed and analyzed all NMR studies; BEM, AC, and SM performed and analyzed all lipidomic studies. WKS, SW, and SPL directed and performed the statistical analyses.

