## Supplementary material for "Sequential Penta-Omic Extraction Method Using Single Biospecimens of Post-mortem Human Brain": Penta-omic Supplemental Figures

**Supplementary Data**

Description of Tables and Figures

1. **Demographics of brain donors, RIN/DV200 values, and RNA/DNA upstream losses:** **Supplementary Table 1**: (**A**) **Demographic** table (**B**) The comparison of RIN values (**C**) The comparison of DV200 values s. (**D**) Amount of RNA and DNA lost in SiMPL-DREx versus the gold standards.

2. **Proteomics and loss in 80% MeOH gold standard extract:** **Supplementary Figure 1**: A Venn diagram of the complementary and different proteins with urea-based extraction of tissue and 80% MeOH **(A)**. A comparison of hydrophobicity **(B)**, pI **(C)** and molecular weight **(D)** of the unique proteins identified in either the MeOH extracted proteins and the unique proteins from both the SiMPL-DREx and Tissue samples.

3. **SiMPL-DREx extraction protocol:** **Supplementary Figure 2**: SiMPL-DREx workflow.

4. **Lipidomics and sequential extractions using the Matyash method:** **Supplementary Figure 3**: (A) The scree plot showing significant variance steadily decreases and the PC2 versus PC3 and PC3 versus PC4 plots. (**B**) The TIC normalized volcano plots of M1 vs M2 (left) and M2 vs M3 showing the FDR significant lipids that are significantly 2-fold different. (**C**) The relative differences in lipid classes extracted in the above comparisons, showing which lipids remain more in subsequent extraction relative to TIC.

5. **Gold standard extraction protocols:** **Supplementary Figure 4**: Gold standard Matyash (left) and 80% MeOH (Right) sequential extraction workflow.

6. **Solvent ratio determination:** **Supplementary Figure 5**: (**A**) $^{1}$H NMR spectra top and table of ratios bottom. NMR Sample preparation, NMR spectral conditions and processing, and calculations describe in Methods. (**B**) Correlation of MTBE/MeOH ratio and water proportion in the aqueous phase. (**C**) Schematic showing the 800 µL and 200 µL taken from the top and bottom phases, respectively. The residual bottom layer was acquired until the top layer reach the bottom and then the remaining top residual layer was taken.

**A**

| **Demographics** | Donor | | | | |
|---|---|---|---|---|---|
| | **1** | **2** | **3** | **4** | **5** |
| Sex | M | F | F | M | F |
| Race | Unk | W | W | B | W |
| Age | 54 | 43 | 57 | 58 | 54 |
| RIN (NBB)* | 7.2 | 6.6 | 6.3 | 7.1 | 6.3 |
| PMI (hr) | 22.4 | 18.0 | 12.0 | 24.9 | 8.7 |

| **Demographics** | Donor | | | | | | | |
|---|---|---|---|---|---|---|---|---|
| | **6** | **7** | **8** | **9** | **10** | **11** | **12** | **13** |
| Sex | M | F | M | M | M | M | F | M |
| Race | W | W | W | W | W | Unk | B | W |
| Age | 53 | 54 | 46 | 43 | 54 | 44 | 28 | 51 |
| RIN (NBB)* | 5.5 | 5.6 | 4.5 | 5.1 | 5.5 | 4.6 | 6.1 | 6.9 |
| PMI (hr) | 17.5 | 19.3 | 18.3 | 21.4 | 18 | 23.3 | 15 | 21.4 |

**B**

| **RIN** | Donor | | | | | | | |
|---|---|---|---|---|---|---|---|---|
| | **6** | **7** | **8** | **9** | **10** | **11** | **12** | **13** |
| SiMPL-DREx Pellet | 4.1 | 1.8 | 1.1 | 6.4 | 3.2 | 2.5 | 3.2 | 2.2 |
| Zymo-Quick Tissue | 3.7 | 1.8 | 1.0 | 6.8 | 2.9 | 2.4 | 2.2 | 1.0 |

**C**

| **DV200** | | | | | | | | | | | | | |
|---|---|---|---|---|---|---|---|---|---|---|---|---|---|
| | **1** | **2** | **3** | **4** | **5** | **6** | **7** | **8** | **9** | **10** | **11** | **12** | **13** |
| SiMPL-DREx Pellet | 82 | 73 | 66 | 84 | 75 | 80 | 57 | 42 | 91 | 77 | 72 | 73 | 66 |
| Zymo-Quick Tissue | 90 | 78 | 76 | 87 | 78 | 73 | 52 | 42 | 88 | 74 | 70 | 71 | 46 |

**D**

| Upstream Loss (%) | SiMPl-DRXx Lipidome | Gold Standard (Matyash) | SiMPl-DRXx Metabolome | Gold Standard (80% MeOH) |
|---|---|---|---|---|
| DNA | 0 | 0.2 ±0.3% | 0 | 2.7 ± 2.5% |
| RNA | 0 | 0 | 0 | 2.0 ± 2.8 % |

**Supplementary Table 1**: (**A**) **Demographic** table of the five donors undergoing -omic analysis (***top***) and eight donors undergoing RIN and DV200 analysis (***bottom***). * RIN values given by NBB. (**B**) The comparison of RIN values between the SiMPL-DREx pellet and Tissue extracted with Zymo-Quick columns. (**C**) The comparison of DV200 values between the SiMPL-DREx pellet and Tissue extracted with Zymo-Quick columns. (**D**) Amount of RNA and DNA lost in SiMPL-DREx versus the gold standards. Nothing was lost in the SiMPL-DREx procedure. Similar to the proteomic results, the 80% MeOH extracts the most RNA and DNA. Very small amount for both, but detectable, and reason for single extraction steps.

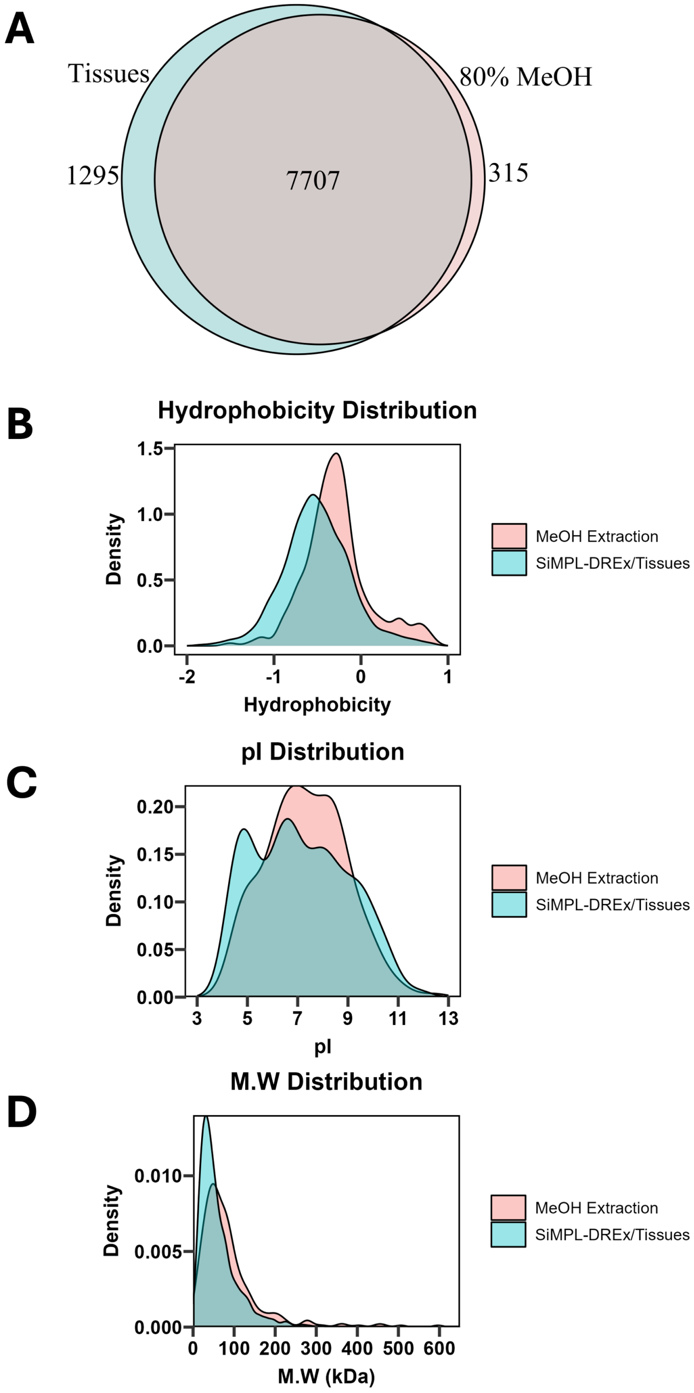


**Supplementary Figure 1**: A Venn diagram of the complementary and different proteins with urea-based extraction of tissue and 80% MeOH **(A)**. A comparison of hydrophobicity **(B)**, pI **(C)** and molecular weight **(D)** of the unique proteins identified in either the MeOH extracted proteins and the unique proteins from both the SiMPL-DREx and Tissue samples. The hydrophobicity was calculated using the peptide package in R with the Kyte-Doolittle approach.

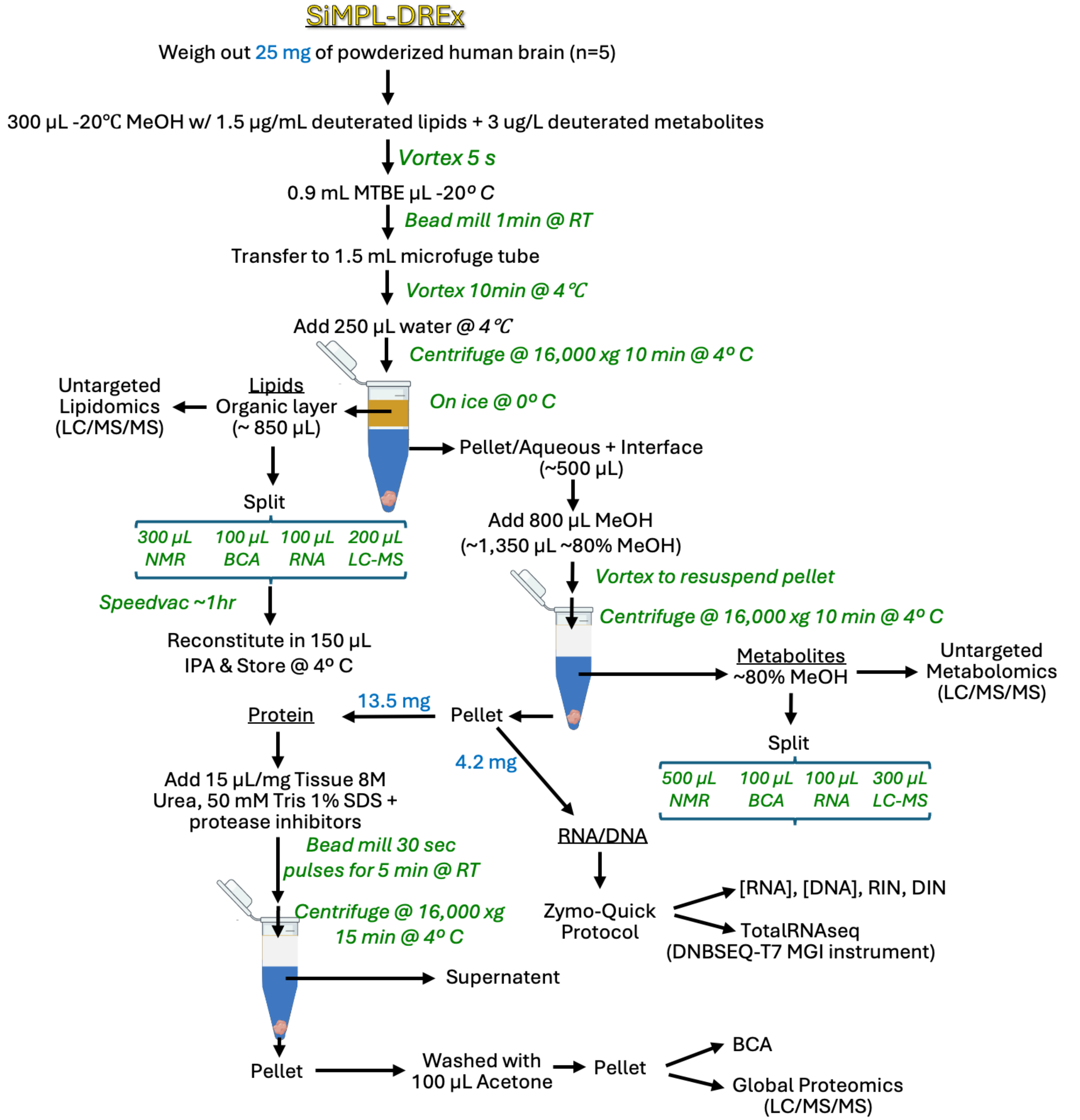


**Supplementary Figure 2**: SiMPL-DREx workflow.

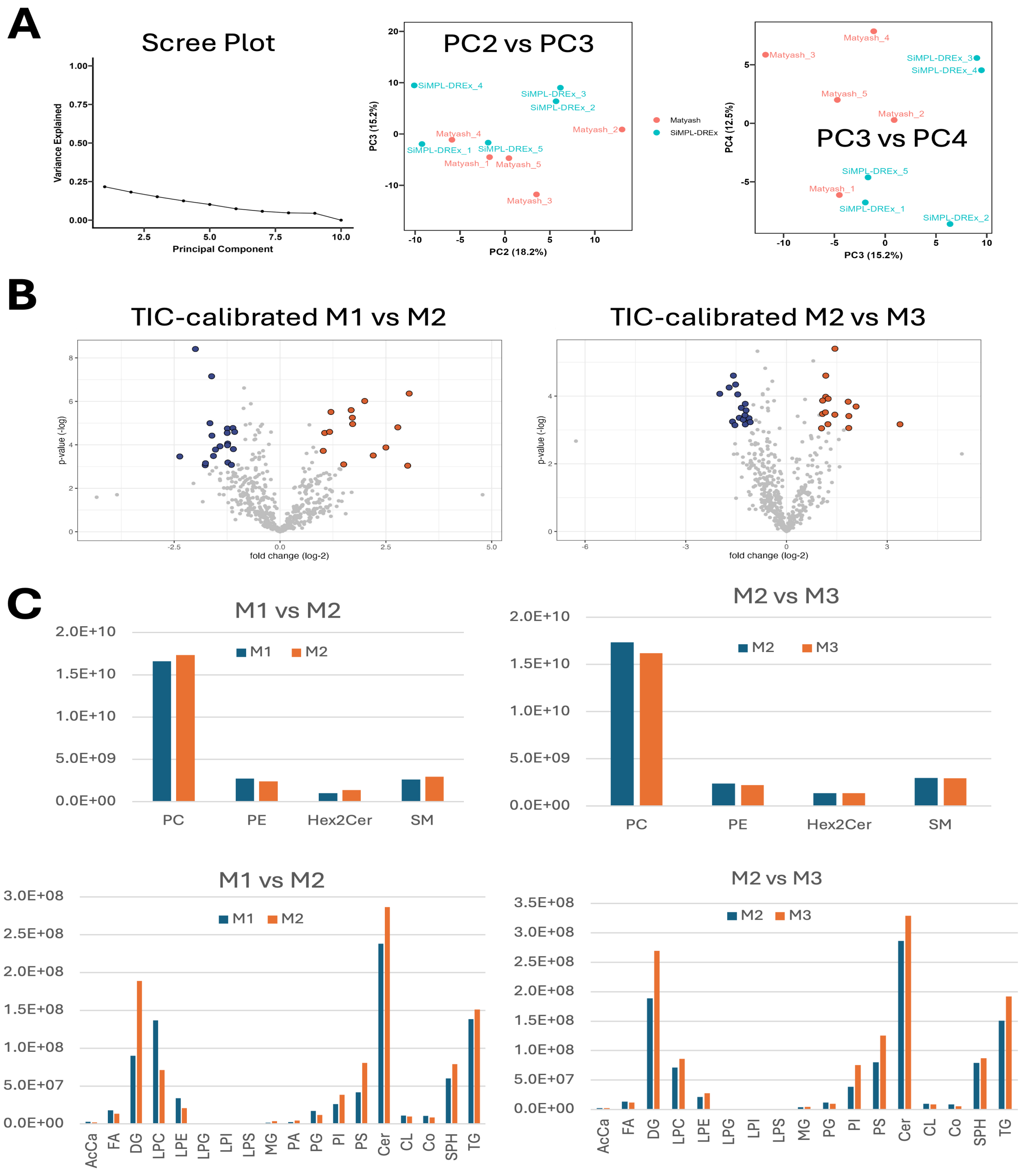


**Supplementary Figure 3**: (**A**) The scree plot showing significant variance steadily decreases and the PC2 versus PC3 and PC3 versus PC4 plots. (**B**) The TIC normalized volcano plots of M1 vs M2 (left) and M2 vs M3 showing the FDR significant lipids that are significantly 2-fold different. (**C**) The relative differences in lipid classes extracted in the above comparisons, showing which lipids remain more in subsequent extraction relative to TIC. None of the comparisons are significantly different but show the trends.

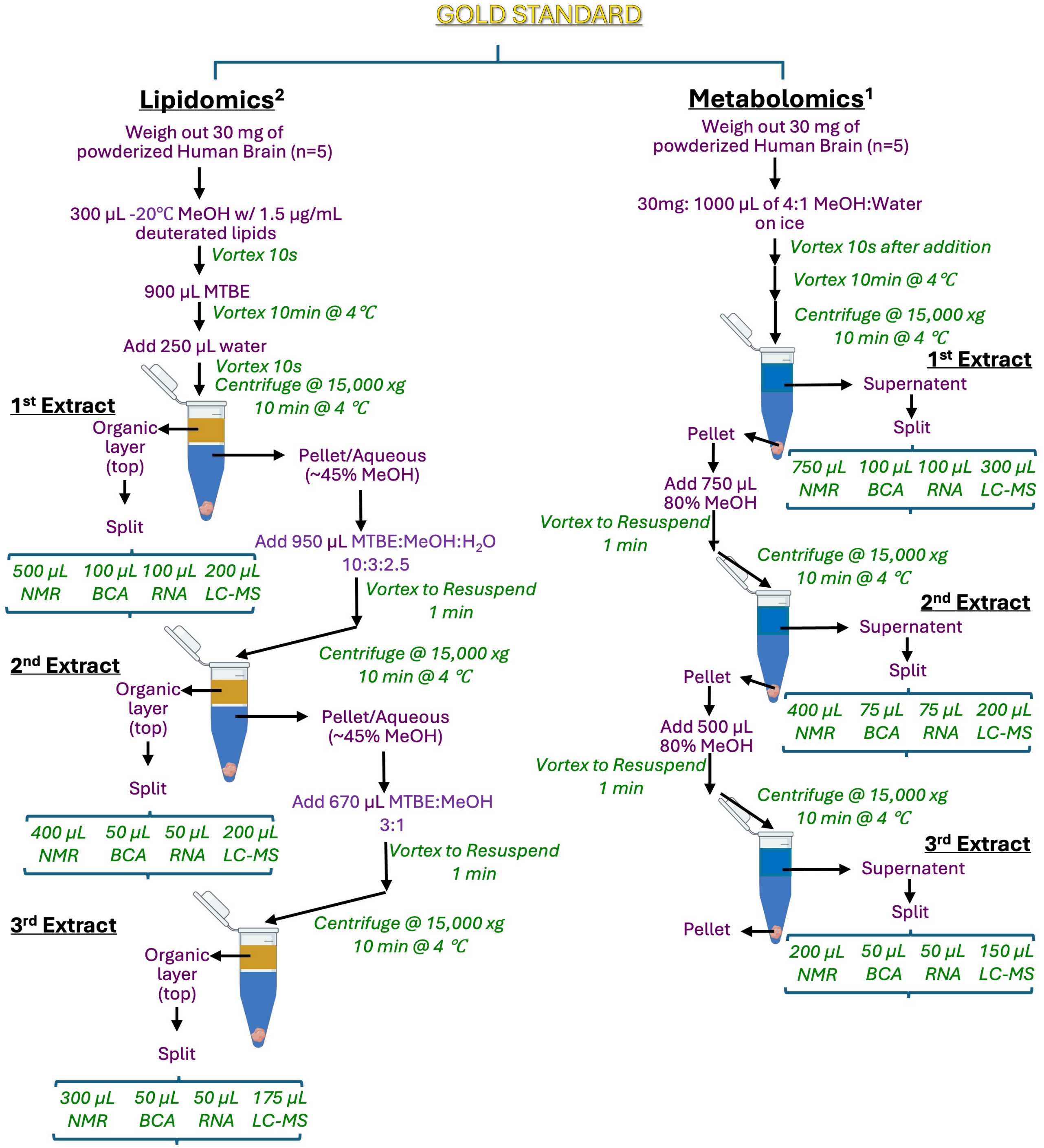


**Supplementary Figure 4**: Gold standard Matyash (left) and 80% MeOH (Right) sequential extraction workflow.

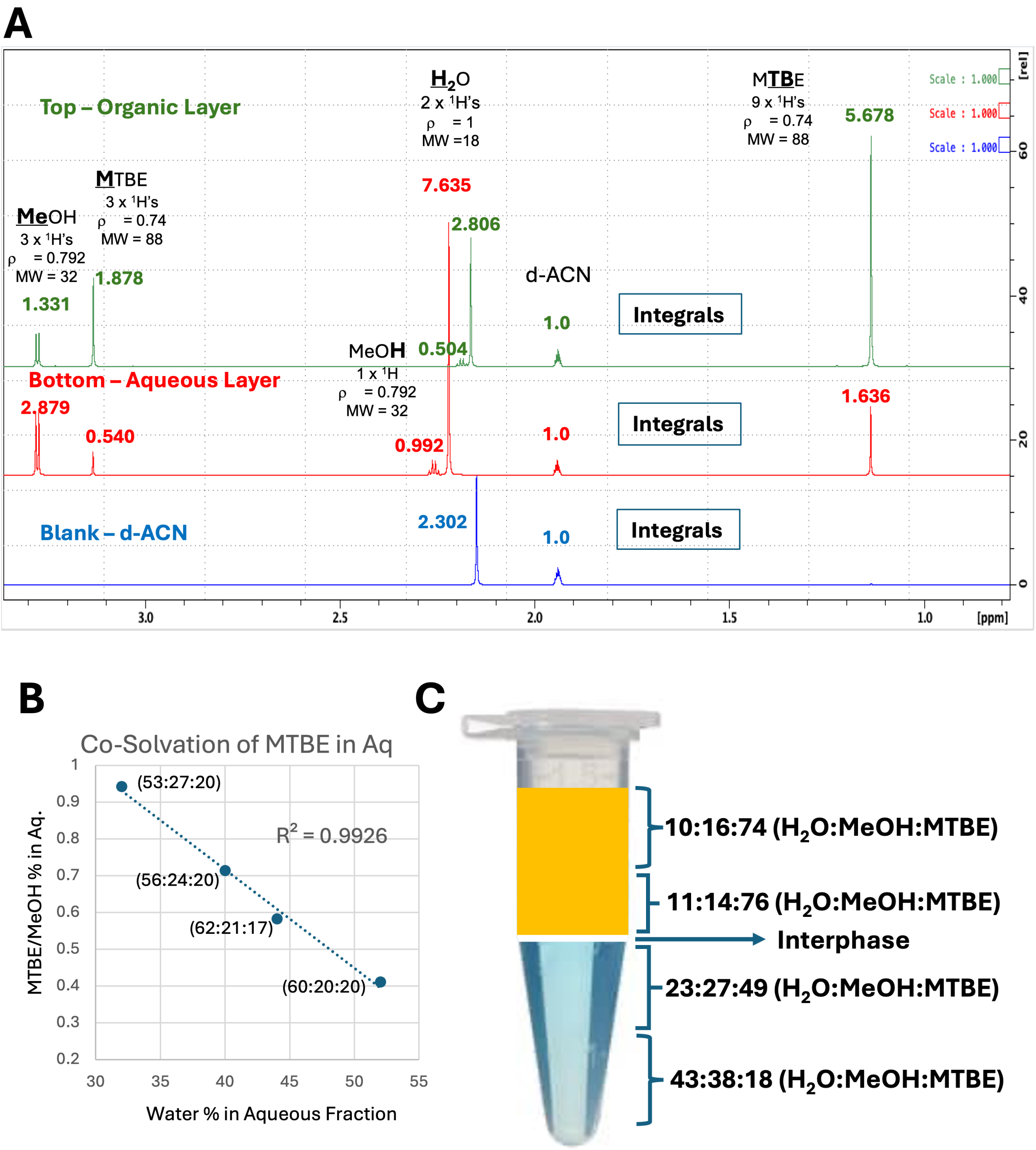


**Supplementary Figure 5**: (**A**) [1]H NMR spectra top and table of ratios bottom. NMR Sample preparation, NMR spectral conditions and processing, and calculations describe in Methods. (**B**) Correlation of MTBE/MeOH ratio and water proportion in the aqueous phase. (**C**) Schematic showing the 800 µL and 200 µL taken from the top and bottom phases, respectively. The residual bottom layer was acquired until the top layer reach the bottom and then the remaining top residual layer was taken.
